# MatP local enrichment delays segregation independently of tetramer formation and septal anchoring in *Vibrio cholerae*

**DOI:** 10.1101/2024.02.08.578905

**Authors:** Elena Espinosa, Jihane Challita, Jean-Michel Desfontaines, Christophe Possoz, Marie-Eve Val, Didier Mazel, Martial Marbouty, Romain Koszul, Elisa Galli, François-Xavier Barre

## Abstract

*Vibrio cholerae* harbours a primary chromosome derived from the monochromosomal ancestor of the Vibrionales (ChrI) and a secondary chromosome derived from a megaplasmid (ChrII). The coordinated segregation of the replication terminus of both chromosomes (*TerI* and *TerII)* determines when and where cell division occurs. ChrI encodes a homolog of *<u>Escherichia coli</u>* MatP, a protein that binds to a DNA motif (*matS*) that is overrepresented in replication termini. Here, we use a combination of deep sequencing and fluorescence microscopy techniques to show that *V. cholerae* MatP structures TerI and TerII into macrodomains, targets them to mid-cell during replication, and delays their segregation, thus supporting that ChrII behaves as a bona fide chromosome. We further show that the extent of the segregation delay mediated by MatP depends on the number and local density of *matS* sites, and is independent of its assembly into tetramers and any interaction with the divisome, in contrast to what has been previously observed in *E. coli*.

## Introduction

Chromosome segregation and cell division must be coordinated in space and time to ensure the faithful transmission of genetic information. In eukaryotes, this is achieved by linking several steps of cytokinesis to the assembly and activity of the mitotic spindle, the eukaryotic chromosome segregation machinery. In bacteria, which lack a functional equivalent of the mitotic spindle, the genomic DNA itself is involved in the regulation of cell division^1–3^. This has been remarkably illustrated in *Vibrio cholerae* - the causative agent of the deadly human diarrhoeal disease of the same name, in which the positioning and timing of assembly of the divisome is primarily dictated by the cellular arrangement and choreography of segregation of the chromosomes^4,5^.

*V. cholerae* is a curved rod-shaped monotrichous bacterium (Fig. 1a). It is intrinsically polarised with a ‘new’ pole resulting from the division of the mother cell and an ‘old’ flagellated pole (Fig. 1a). Its genome is distributed on a primary chromosome derived from the monochromosomal ancestor of the Vibrionales (ChrI) and a secondary chromosome derived from a megaplasmid (ChrII)^6^. ChrI and ChrII both carry a single origin of replication, *ori1* and *ori2*, with replication terminating in a diametrically opposite zone, *TerI* and *TerII*, which contains a site dedicated to the resolution of chromosome dimers by the XerCD/FtsK machinery, *dif1* and *dif2*, respectively (Fig. 1a and 1b)^7,8^. Both are arranged longitudinally in the cell, with *TerI* and *TerII* migrating from the new pole to mid-cell in the course of replication (Fig. 1a)^9,10^. Binding sites for a DNA-associated protein that inhibits divisome assembly, SlmA, are enriched all along ChrI and ChrII except for *TerI* and *TerII* (Fig. 1a)^1,4^. Thus, the positioning of *TerI* and *TerII* restricts septum formation to mid-cell at the end of the cell cycle (Fig. 1a)^4,5^. Accordingly, we observed cell division defects in MCH1, a synthetic monochromosomal strain in which all of ChrII except *ori2* and its partition machinery is inserted into ChrI at the *dif1* locus^4^.

**Fig. 1.**
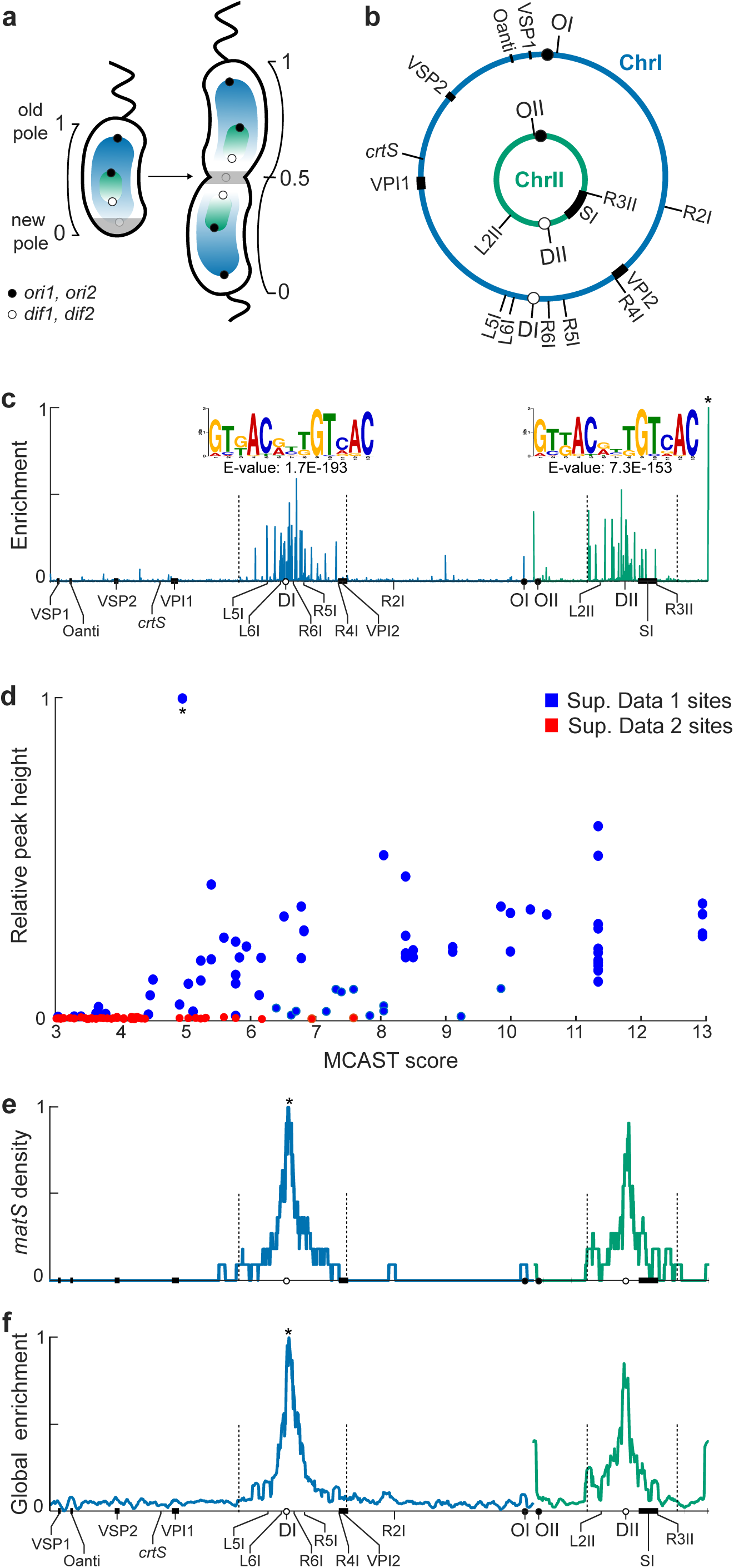
Numerous *matS* sites flank both *dif1* and *dif2*. **a** Schematic of newborn and dividing *V. cholerae* cells. Spiral line: flagellum; grey shading: cell division proteins; blue and green gradients: SlmA-mediated nucleoid occlusion gradient; black shaded circles: *ori1* and *ori2*; white shaded circles: *dif1* and *dif2*. By convention, the pole resulting from the mother cell division is referred to as the new pole and the opposite pole, where the flagellum is located, is referred to as the old pole. **b** *V. cholerae* chromosome maps. ChrI and ChrII are depicted in blue and green, respectively. Black rectangles: super integron (SI), Vibrio Pathogenicity Islands (VPI1, VPI2), Vibrio Seventh pandemic Islands (VSP1, VSP2) and O-antigen cluster (Oanti), as indicated. The position of *crtS* and of the loci tagged for microscopic observation are indicated. **c** ChIP-seq analysis of the binding of *V. cholerae* MatP. Blue and green lines: relative sequence enrichment profiles along ChrI and ChrII, respectively. The profiles were smoothed using a sliding average distance of 100 bp and scaled between 0 (lowest enrichment locus along the whole genome) and 1 (highest enrichment locus over the whole genome, indicated with a star). The MEME logos of the *matS* sites found on ChrI and ChrII are shown above their respective profiles. The position weight matrix of the ChrII sites matches that of the ChrI sites (Two-sided Pearson correlation, p-value: 1.95E-11, q-value: 3.89E-11). Genomic features of interest are displayed on the x-axis. Black dotted lines indicate the boundaries of the *TerI* and *TerII* MDs. **d** Relative enrichment of the sequences as a function of their MCAST score. Blue and Red: N16961 *V. cholerae matS* sites bound and not bound by MatP, respectively. A star indicates the motif showing the highest enrichment. **e** 40 kbp sliding average profile of the number of *matS* sites along ChrI and ChrII indicating the expected enrichment of MatP-bound sequences at the centre of *TerI* and *TerII* induced by the local density of *matS* sites. **f** 40 kbp sliding average profile of Fig. 1c ChIP-seq data, showing the actual enrichment of MatP at the centre of *TerI* and *TerII*. Genomic features of interest are displayed on the x-axis.

But what is the mechanism that drives the choreography of *TerI* and *TerII* segregation? ChrI encodes for a homolog of MatP, a protein that binds as a dimer to a DNA motif, *matS*, that is overrepresented in the replication terminus region of the *Escherichia coli* chromosome, *Ter* ^11,12^. MatP plays two roles in *E. coli*: (i) it structures *Ter* into a macrodomain (MD) by limiting long-range *cis* contacts (*cis* contacts between DNA sequences separated by more than 280 kbp) between *Ter* and the rest of the genome^13,14^ and (ii) it ensures the colocalisation of sister *Ter* copies at mid-cell until the onset of septation^15^. The last 20 carboxy-terminal amino acid residues of MatP induce the formation of tetramers (dimers of dimers) that can link *matS* sites to each other and/or to other DNA segments^12,16^. Tetramers are dispensable for MD formation^14^. Their role in the initial post-replicative pairing of sister *Ter* copies is still under debate^16,17^. However, the prolonged positioning of sister *Ter* copies at mid-cell is thought to be due to the interaction of the tetramerization domain with a cell division protein, ZapB^15,16,18,19^. Functional homologs of MatP and ZapB similarly link the *Ter* of the *Caulobacter crescentus* chromosom*e* to the divisome^20,21^. Based on the *E. coli matS* motif consensus, 34 and 14 putative *matS* sites have been identified in *TerI* and *TerII*, respectively^13^. Early work showed that the deletion of *matP* led to the early separation of sister *dif1* loci and the erratic positioning of *dif2* loci^10^, and impaired cell division^4^. However, sister *dif1* loci were found to remain associated independently of divisome assembly during the recovery of *V. cholerae* spheroplasts to a proliferative rod-shaped state, contradicting the idea that MatP prolonged the colocalization of sister *TerI* copies by tethering them to the divisome^22^. In addition, sister *dif2* loci were found to separate and move away from mid-cell prior to sister *dif1* copies at the onset of constriction^10^. These observations questioned the role and action mode of MatP in *V. cholerae*.

Here, we have used a combination of deep sequencing and fluorescence microscopy techniques to investigate the role of MatP in *V. cholerae*. We show that it binds to a very large number of sites in *TerI* and *TerII*, similar to but more diverse than the *E. coli matS* sites. We also show that MatP structures *TerI* and *TerII* into macrodomains, targets them to mid-cell in the course of replication and delays their segregation, demonstrating that ChrII is organised and behaves as a *bona fide* chromosome. However, in contrast to what has been observed in *E. coli*, we show that *V. cholerae* MatP can delay segregation independently of tetramer formation and any divisome interaction, and that the extent of the segregation delay is determined by the number and local density of *matS* sites.

## Results

### Numerous *matS* sites flank both *dif1* and *dif2*

Based on the *E. coli matS* motif, 14 putative *matS* sites were identified in *TerII*^13^. This was in apparent contradiction with the early separation and migration away from mid-cell of sister *TerII* copies all the more as two *matS* sites are sufficient to bring a plasmid to mid-cell in *E. coli*^9,10,15^. However, we noticed that a conserved amino acid motif differentiated the Vibrionales and Enterobacterales MatP proteins^6^. It raised the possibility that *V. cholerae* MatP bound to different DNA sites than *E. coli* MatP, which would be under-represented in *TerII*. To check this possibility, we used ChIP-seq to explore the binding of *V. cholerae* MatP on ChrI and ChrII *in vivo* (Methods). To this end, we introduced a FLAG tag allele of *matP* in place of the *matP* gene (*matP3flag*). The choreography of segregation of sister *dif1* copies in cells harbouring the *matP3flag* allele was identical to that of *matP^+^*cells, in contrast to Δ*matP* cells, indicating that the allele was functional (Supplementary Fig. 1a).

The *matP3flag* ChIP-seq profile showed clearly distinguishable peaks (Fig. 1c). No peaks were observed in the ChIP-seq profile obtained in a strain carrying a FLAG version of a C-terminal truncation allele of *matP* that, based on the homology with *E. coli matP*, was expected to prevent DNA binding by abolishing dimer formation (Supplementary Fig. 1b). These results indicated that the peaks of the *matP3flag* ChIP-seq profile corresponded to MatP-binding sites and that the C-terminal truncation allele of *V. cholerae matP* prevented the formation of MatP dimers, i. e. it was a *matP* Δ*dimer* allele.

We identified 43 and 35 *matS* sites on ChrI and ChrII, respectively (Supplementary Data 1, Supplementary Fig. 1c). Their consensus sequence was similar to the *E. coli matS* motif (Supplementary Fig. 1d). No significant difference was observed in the sequence logos of the ChrI and ChrII *matS* sites (Fig. 1c). We detected 66 additional putative *matS* sites in the *V. cholerae* genome based on the *V. cholerae matS* consensus (Supplementary Data 2). No enrichment was observed at the location of these sites, suggesting that they were not functional (Supplementary Fig. 2).

Most of the *V. cholerae matS* sites were located in the terminal proximal part of the chromosomes (Fig. 1c, Supplementary Data 1). The only notable exception was a site located in the immediate vicinity of *ori2*. We previously noticed that *TerI* harboured a very strong SlmA binding site (SBS) and a binding site for ParAB2 (*parS2*) separated by only 68 bp^4,23^. Therefore, we suspect that the presence of a *matS* site near *ori2* was caused by a translocation event between this region and *TerI*.

The relative height of the ChIP-seq peaks tended to be higher near *dif1* and *dif2* (Fig. 1c). This was not explained by any *matS* sequence specificity (Fig. 1d), but correlated with the local density of *matS* sites (Fig. 1e). As the density of *matS* sites increases in the vicinity of *dif1* and *dif2*, the cumulative binding of MatP was the highest at *dif1* and *dif2* (Fig. 1f).

### MatP structures *TerI* and *TerII* into macrodomains

The *Ter* MD of the *E. coli* chromosome was defined as the longest stretch of DNA where *matS* sites are separated by less than 100 kbp^13^. Based on the same criteria, 40 out of the 43 ChrI *matS* sites defined a 659 kbp MD in the *TerI* zone, and 34 out of the 35 ChrII *matS* sites defined a 546 kbp MD in the *TerII* zone (Fig. 1c, Supplementary Data 1). The mean density of *matS* sites within these zones (∼6 sites per 100 kbp) was twice higher than the mean density of *matS* sites in the *E. coli Ter* MD (∼3 sites per 100 kbp^13^). To check that MatP structured *TerI* and *TerII* into MDs, we applied 3C coupled with deep sequencing (3C-seq) to exponentially growing *matP^+^* and Δ*matP* N16961 cells. The contact map of WT cells showed a single strong diagonal resulting from the enrichment of contacts between adjacent loci of each chromosome (Supplementary Fig. 3a)^24^. The width of the diagonal was smaller in the *TerI* and *TerII* zones than in the rest of the genome, reflecting a reduction of long-range *cis* contacts (Supplementary Fig. 3a)^24^. Long-range *cis* contacts were recovered in Δ*matP* cells (Fig. 2, Supplementary Fig. 3b). In addition, the deletion of *matP* increased contacts between *ori2* and *TerII*, suggesting that ChrII had entirely lost its structural conformation (Fig. 2).

**Fig. 2.**
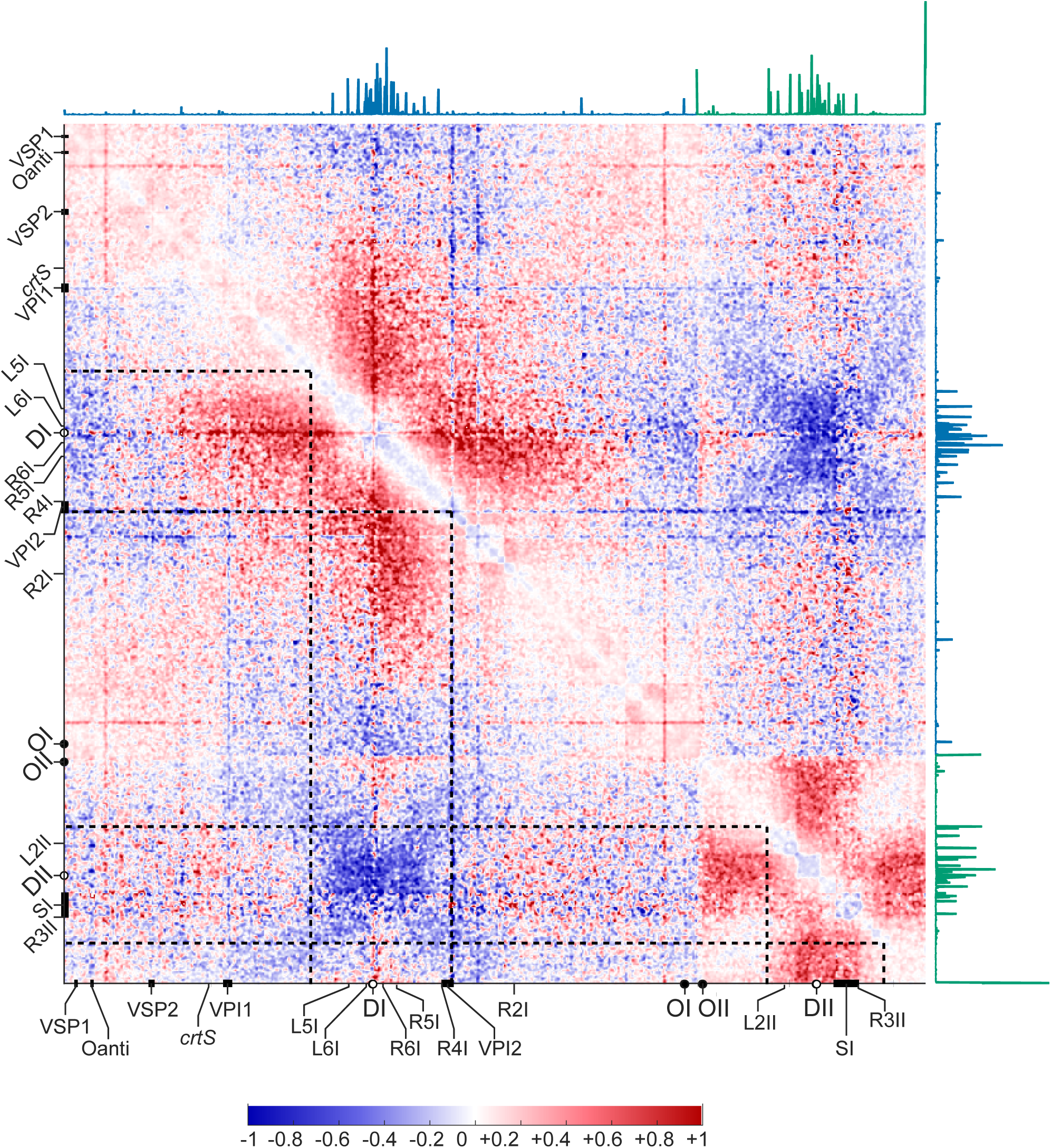
MatP structures *TerI* and *TerII* into macrodomains. Ratio plot of the 3C-seq maps of Δ*matP* and *matP ^+^* cells. A decrease or increase in contacts in 5-kbp bins in Δ*matP* cells compared with *matP ^+^* cells is represented with a blue or red colour, respectively. White indicates no differences between the two conditions. Genomic features of interest are indicated on the bottom and left axes. The ChIP-seq profile obtained with the *matP3flag* allele is shown on the top and right axes. Black dotted lines indicate the boundaries of the *TerI* and *TerII* MDs.

### Deletion of *matP* leads to erratic positioning of *TerI* and *TerII*

We monitored the position of 12 loci in fluorescence snapshot microscopy images (Methods). They included a locus at 53 kbp from *ori1* (OI), a locus at 24 kbp from *ori2* (OII), a locus right next to *dif1* (DI) and a locus at 10 kbp from *dif2* (DII), 4 loci within the *TerI* MD (L5I, L6I, R6I and R5I), 2 loci on the right replication arm of ChrI (R2I and R4I), and one locus on each of the two replication arms of ChrII (L2II and R3II) (Fig. 1b).

Demographs (Methods) showed that the deletion of *matP* altered the positioning and choreography of segregation of all loci but OI and OII (Supplementary Fig. 4). To quantify the effect of the deletion of *matP*, we analysed the cellular position of spots before their duplication in cells at an early stage of the cell cycle and after their duplication in cells at a late stage of the cell cycle, hereafter referred to as young and old cells (Methods). In young *matP^+^*cells, *TerI* loci were all constrained at a median distance of ∼15% of the cell length from the new pole (Fig. 3a). The median distance increased to ∼33% of the cell length in Δ*matP* cells (Fig. 3a). In contrast, *TerII* loci were positioned at a median distance of ∼30% of the cell length in both *matP^+^* and Δ*matP* cells (Fig. 3a). However, the deletion of *matP* increased the interquartile distribution of the distance of *TerII* loci from the new pole to ∼27% of the cell length, as it did for *TerI* loci (Fig. 3a). The same trend was observed for the position of sister spots of old cells (Supplementary Fig. 5a).

**Fig. 3.**
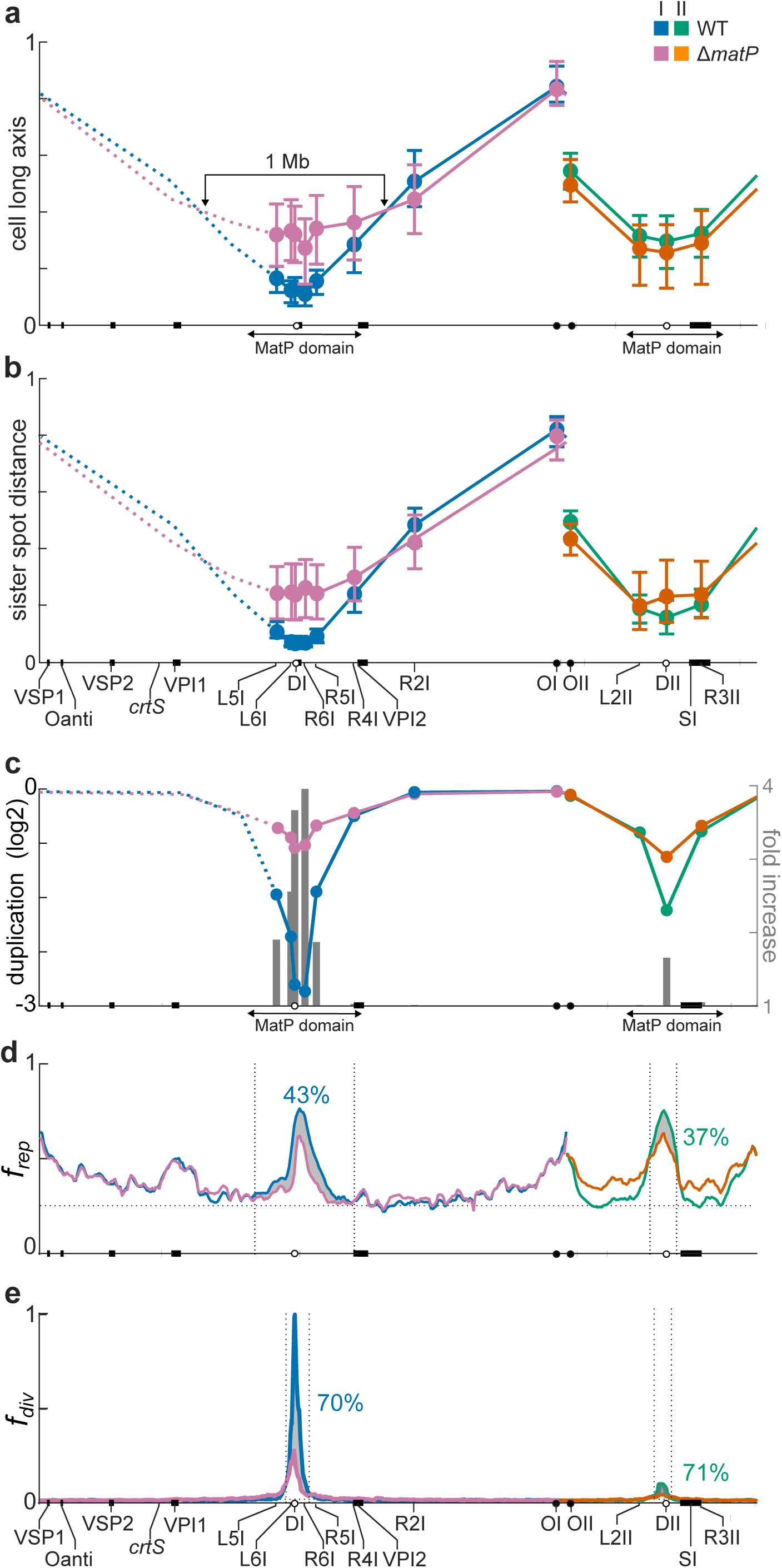
MatP increases cohesion in the immediate vicinity of *dif1* and *dif2*. **a** Relative longitudinal axis position of single spots. The new and old pole positions are set to 0 and 1, respectively. Results correspond to a median number of 591 cells for each locus (Supplementary Data 3). **b** Relative distance along the longitudinal axis between sister copies of each tagged locus in old cells. Results correspond to a median number of 319 cells for each locus (Supplementary Data 3). The median (disk), first and third quartile (horizontal marks) of the positions/sister distances are shown for each tagged locus. Solid lines: expected position/sister distance of intermediate loci; Dashed lines: the expected symmetrical position/sister distance of loci on the left replication arm of ChrI; Blue and pink: ChrI; green and orange: ChrII; blue and green: *matP ^+^*cells (WT); pink and orange: Δ*matP* cells (Δ*matP*); Arrows: *TerI* and *TerII* MDs. Genomic features of interest are displayed on the x-axis. **c** log2 of the spot duplication frequency at each locus (blue, pink, green and orange lines) and increase in duplication frequency at each locus in Δ*matP* cells (grey bars). **d** Cre/*loxP*-based Hi-SC2 analysis of the relative frequency of contacts of newly-replicated copies behind replication forks. **e** FtsK/XerCD/*dif1*-based Hi-SC2 analysis of the relative frequency of contacts of sister copies during septum closure. Results are shown at a 40 kbp resolution. Black dotted lines indicate the zones where sister chromatid contacts are less frequent in Δ*matP* cells than in WT cells. Grey surfaces highlight the global reduction of sister chromatid contacts in these zones. Genomic features of interest are displayed on the x-axis. The corresponding percentages of reduction is indicated. Hi-SC2 analysis replicates are shown in Supplementary Fig. 5b,c. Source data of panel a, b and c are provided as a Source Data file.

To further highlight the effect of MatP on the cellular arrangement of ChrI and ChrII, we plotted the median distance between sister spots in old cells (Fig. 3b). In *matP* ^+^ cells, DI, L5I, L6I, R6I, R5I and R4I sister spots were closer than expected from a linear increase as a function of their genomic distance from OI (Fig. 3b). In particular, the median distance between DI, L6I, and R6I sister spots was lower than ∼4% of the cell length (Fig. 3b). Deletion of *matP* increased the median distance between sister spots of *TerI* loci to ∼27% of the cell length (Fig. 3b). In contrast, DII, L2II and R3II sister spots were at a median distance from each other of ∼15% of the cell length in both *matP^+^* and Δ*matP* cells (Fig. 3b). However, the deletion of *matP* increased the interquartile distribution of *TerII* sister spot distances to ∼25% of the cell length, as it did for *TerI* loci (Fig. 3b). The deletion of *matP* also altered R4I and R2I sister spot distances, notably by increasing their interquartile distribution (Fig. 3b).

Taken together, these results showed that MatP constrained the central part of the *TerI* MD to the new pole in young cells and to mid-cell in old cells, which affected a large 1 Mbp zone extending outside of the *TerI* MD (Fig. 3a and 3b). In contrast, MatP was not able to tether *TerII* loci at the new pole in young cells and at mid-cell in old cells. However, our results suggested that the positioning of all ChrI and ChrII loci but OI and OII was more erratic in Δ*matP* cells.

### MatP is involved in *dif1*- and *dif2*-cohesion at the onset of division

In *matP^+^* cells, the spot duplication frequency of ChrI loci did not follow their timing of replication^7^: it was similar at OI and R2I, but dropped abruptly inside the *TerI* MD (Fig. 3c). Likewise, the spot duplication frequency of DII was lower than expected from its replication timing (Fig. 3c). Deletion of *matP* suppressed the separation delay of *TerI* loci and that of DII, with the spot duplication frequency now reflecting their timing of replication (Fig. 3c). In addition, the duplication increase was much larger in the immediate vicinity of DI and DII than at loci at the border of the *TerI* and *TerII* MDs (grey bars in Fig. 3c). These observations suggested that MatP increased cohesion.

To further investigate the role of MatP in sister chromatid cohesion, we monitored the frequency of sister chromatid contacts behind replication forks (*f*_*ref*_) and at the time of cell division (*f*_*div*_) with a high-resolution whole-genome method, Hi-SC2 (Methods).

In *matP* ^+^ strains, *f*_*ref*_ was higher in the *TerI* and *TerII* MDs than in the rest of the genome and was maximal at *dif1* and *dif2* (Fig. 3d). Deletion of *matP* did not modify *f*_*ref*_along ChrI except in the *TerI* MD where ∼40% of the sister chromatid contacts were lost (Fig. 3d, Supplementary Fig. 5b). It increased *f*_*ref*_ in the replication arms of ChrII except in a 150 kbp zone within the *TerII* MD where ∼40% of the sister chromatid contacts were lost (Fig. 3d, Supplementary Fig. 5b).

In *matP^+^* cells, *f*_*div*_ was null at all genomic positions except in the immediate vicinity of *dif1* and *dif2* (Fig. 3e, Supplementary Fig. 5c). Note that a large *dif1* cassette, which allows intra-molecular recombination, was excised at a frequency greater than 50% in a third of ChrI and half of ChrII (Supplementary Fig. 5d), excluding the idea that Xer recombination could only occur in the vicinity of *dif1* and *dif2* because of FtsK is directed towards them by orienting polar sequences ^8,25–28^. In Δ*matP* cells, ∼70% and 60% of *TerI* and *TerII* sister chromatid contacts at cell division were lost, respectively (Fig. 3e, Supplementary Fig. 5c).

Taken together, these results indicated that MatP was involved in the post-replicative cohesion of loci within the *TerI* and *TerII* MDs and played a major role in the persistence of cohesion until the initiation of cell division in the immediate vicinity of *dif1* and *dif2*.

### MatP extends cohesion between consecutive MatP domains

To gain further insight into the role of MatP in the arrangement and segregation of chromosomes, we analysed its action in the synthetic monochromosomal MCH1 strain (Fig. 4a)^29^. Previous work showed that replication of the MCH1 chromosome starts at *ori1* and ends in *TerII*^4^, and that chromosome dimers are efficiently resolved by Xer recombination at *dif2*^29^. ChIP-seq showed that there are 3 MatP domains in the MCH1 chromosome, the *TerII* MD and the Right and Left part of the *TerI* MD (Fig. 4b, Supplementary Fig. 6a, Supplementary Data 4). The most highly enriched sequences were located at the centre of the *TerII* MD (Fig. 4b, note that the highly enriched *matS* site located near *ori2* in N16961 is absent from the MCH1 chromosome). Sequences in the Right *TerI* MD were as enriched as sequences in the *TerII* MD, but the enrichment of sequences in the Left *TerI* MD was very poor (Fig. 4b, Supplementary Fig. 6a).

**Fig. 4.**
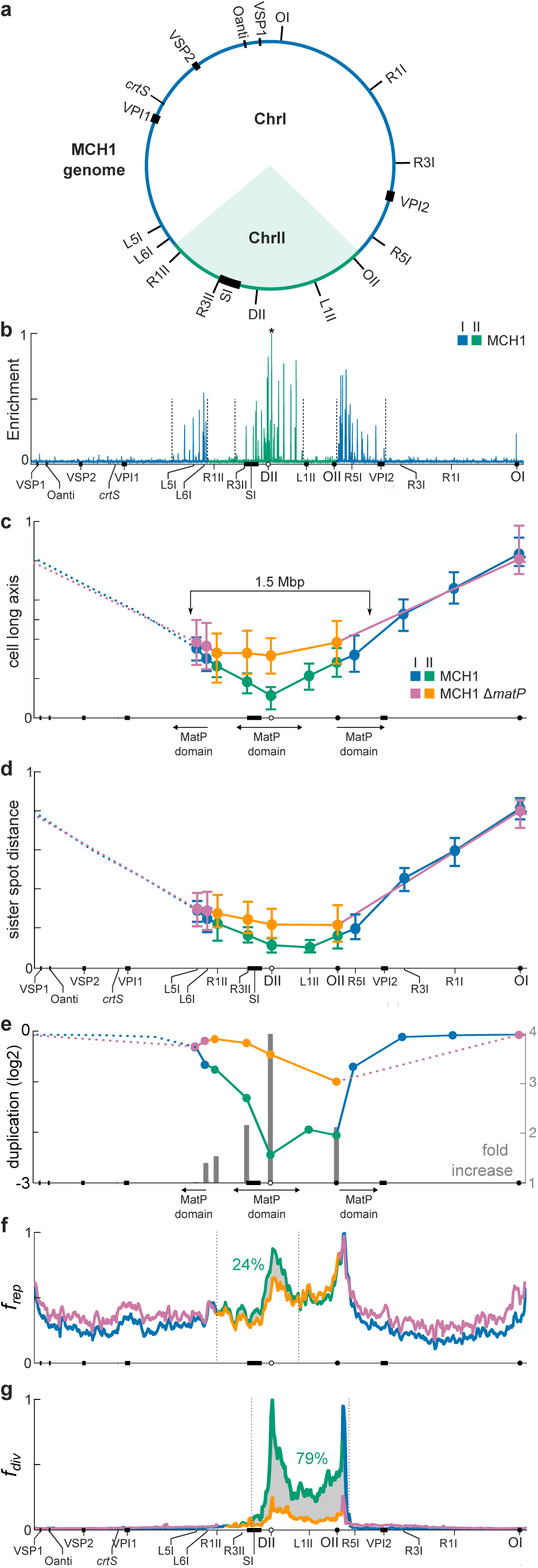
MatP increases cohesion in highly enriched *matS* regions. **a** MCH1 chromosome map. Regions originating from ChrI and ChrII are depicted in blue and green, respectively. *ori1*, *dif2*, SI, VPI1, VPI2, VSP1, VSP2, O-antigen cluster, *crtS* and positions of the loci tagged for microscopic observation are indicated as in Fig. 1. **b** ChIP-seq analysis of the binding of *V. cholerae* MatP in MCH1. Relative sequence enrichment profile along MCH1 genome (100 bp sliding average). The profile was scaled between 0 (lowest enrichment locus along the whole genome) and 1 (highest enrichment locus over the whole genome, indicated with a star). Genomic features of interest are displayed on the x-axis. **c** Relative longitudinal axis position of each tagged locus in young cells. Results correspond to a median number of 638 cells for each locus. **d** Relative distance along the longitudinal axis of sister copies of each tagged locus in old cells. Results correspond to a median number of 417 cells for each locus (Supplementary Data 3). The median (disk), first and third quartile (horizontal marks) of the positions/sister distances are shown for each tagged locus. Solid lines: expected position/sister distance of intermediate loci; Dashed lines: the expected symmetrical position/sister distance of loci on the left replication arm; Blue and pink: regions originating from ChrI; green and orange: regions originating from ChrII; blue and green: *matP^+^* cells (MCH1); pink and orange: Δ*matP* cells (MCH1 Δ*matP*); Arrows: limits of Left, Ter and Right MDs. Genomic features of interest are displayed on the x-axis. **e** log2 of the spot duplication frequency at each locus (blue, pink, green and orange lines) and increase in duplication frequency at each locus in MCH1 Δ*matP* cells (grey bars). **f** Cre/*loxP*-based Hi-SC2 analysis of the relative frequency of contacts of newly-replicated copies behind replication forks. **g** FtsK/XerCD/*dif1*-based Hi-SC2 analysis of the relative frequency of contacts of sister copies during septum closure. Results are shown at a 40 kbp resolution. Black dotted lines indicate the zones where sister chromatid contacts are less frequent in MCH1 Δ*matP* cells than in MCH1 cells. Grey surfaces highlight the global reduction of sister chromatid contacts in these zones. Genomic features of interest are displayed on the x-axis. The corresponding percentage of reduction is indicated. Hi-SC2 analysis replicates are shown in Supplementary Fig. 6e and 6f. Source data of panels c, d and e are provided as a Source Data file.

We monitored the position of 11 loci in fluorescence snapshot microscopy images, including the OI, OII and DII loci, 5 loci on the right replication arm (R1I, R3I, R5I, OII and L1II) and 4 loci on the left replication arm (R3II, R1II, L6I and L5I) of the chromosome (Fig. 4a). Demographs showed that MatP was involved in the choreography of segregation of L6I, R3II, DII, L1II and OII (Supplementary Fig. 6b and 6c). In young *matP* ^+^ MCH1 cells, the distance between the spot of each labelled locus and the new pole increased almost linearly as a function of their genomic distance from DII (Fig. 4c). DII was closer to the new pole than it was in young N16961 cells (Fig. 4c). The same trend was observed in old cells for the distance of sister spots from the centre of the cell (Supplementary Fig. 6d) and the distance between sister spots (Fig. 4d). Loci comprised between OII and L6I were further away from the new pole in young MCH1 Δ*matP* cells than in *matP^+^* cells (Fig. 4c). DII was at the same distance from the new pole in young MCH1 Δ*matP* cells than it was in N16961 Δ*matP* cells (Fig. 4d). The same trend was observed for the distance of sister spots from the centre of the cell (Supplementary Fig. 6d) and the distance between sister spots (Fig. 4d). These results suggested that MatP played a significant role on the arrangement of the MCH1 chromosome.

Strikingly, the spot duplication frequency was low in the vicinity of DII but also at OII, L6I, R1II, R3II, R5I and L5I (Fig. 4e). Correspondingly, *f*_*ref*_was high within *TerII*, the right half of *TerI*, and all the intermediate positions between these two zones (Fig. 4f, Supplementary Fig. 6e). In addition, *f*_*div*_ was highly elevated in the 800 kbp zone comprised between DII and R5I (Fig. 4g, Supplementary Fig. 6f). Deletion of *matP* only slightly increased *f*_*ref*_ in the regions corresponding to ChrI (Fig. 4f, Supplementary Fig. 6e). However, it led to a ∼25% reduction in sister chromatid contacts behind replication forks in the 150 kbp zone surrounding *dif2* and resulted in the loss of ∼80% of the sister chromatid contacts in the DII-R5I region at the time of cell division (Fig. 4g, Supplementary Fig. 6f). These results indicated that MatP was able to maintain together sister copies of loci between the MCH1 *TerII* MD and the Right *TerI* MD, but failed to maintain together sister copies of loci in the Left *TerI* MD.

### MatP-mediated cohesion depends on the number and local density of *matS* sites

The number and density of *matS* sites of the Left *TerI* MD of the MCH1 chromosome is lower than that of the Right *TerI* MD, which suggested that a threshold had to be reached for the effect of MatP on cohesion to be visible (Fig. 4b). To test this hypothesis, we displaced two *matS* sites adjacent to *dif1* in ChrI next *dif2* in ChrII (Fig. 5a, JVV024). It decreased the density of *matS* sites in the *TerI* MD from 6.1 to 5.8 sites per 100 kbp and increased the density of *matS* sites in the *TerII* MD from 6.2 to 6.6 sites per 100 kbp (Supplementary Data 5), which mitigated the difference in the enrichment of MatP at DI and DII (Fig. 5b). Shifting the two *matS* sites increased *f*_*div*_ at the DII locus (Fig. 5c, Supplementary Fig. 7a). Demographs showed no obvious difference between the choreography of segregation of DI and DII in JVV024 and N16961 (Supplementary Fig. 7b). However, the distance separating DI sister copies was slightly but significantly higher in JVV024 cells than in N16961 cells (Fig. 5d, Supplementary Table 1). In addition, the relative timing of appearance of two DI spots loci was now similar to that of DII spots (Supplementary Fig. 7c). Correspondingly, the proportion of cells showing a single DII spot and two DI spots was significantly higher than that of cells showing two DII spots and a single DI spot in JVV024 compared to N16961 (Fig. 5e, Supplementary Table 2). These results suggested that MatP influenced cohesion at the time of cell division at positions where it was highly enriched, namely *dif1* and *dif2* in N16961 (Fig. 1f), and *dif2* and the junction of ChrII DNA with the Right *TerI* MD in MCH1 (Supplementary Fig. 6a). To test this hypothesis, we constructed a strain in which the right part of the *TerI* MD was inverted, EPV487 (Fig. 5f). The inversion created a cleft in the MatP ChIP-seq profile at *dif1* (Fig. 5g, Supplementary Data 6). The position with the highest MatP enrichment was now at the junction of the inverted fragment and the right replication arm (Fig. 5g). The *f*_*div*_ profile followed the local enrichment of MatP (Fig. 5h, Supplementary Fig. 7d). As with N16961 and MCH1, the elevated *f*_*div*_ was mainly associated with MatP (Supplementary Fig. 7e). Demographs showed no obvious difference between the choreography of segregation of DI and DII in EPV487 and N16961 (Supplementary Fig. 7b). However, the inversion significantly affected the distance between DI and DII sister copies (Fig. 5d, Supplementary Table 1). In addition, the relative timing of appearance of two DI spots was now earlier than that of DII spots (Supplementary Fig. 7c). Correspondingly, the proportion of cells showing two DI spots and a single DII spot was significantly higher than that of cells showing a single DI spot and two DII spots in EPV487 cells compared to N16961 cells, reflecting the decrease in cohesiveness of the DI sister copies (Fig. 5e, Supplementary Table 2).

**Fig. 5.**
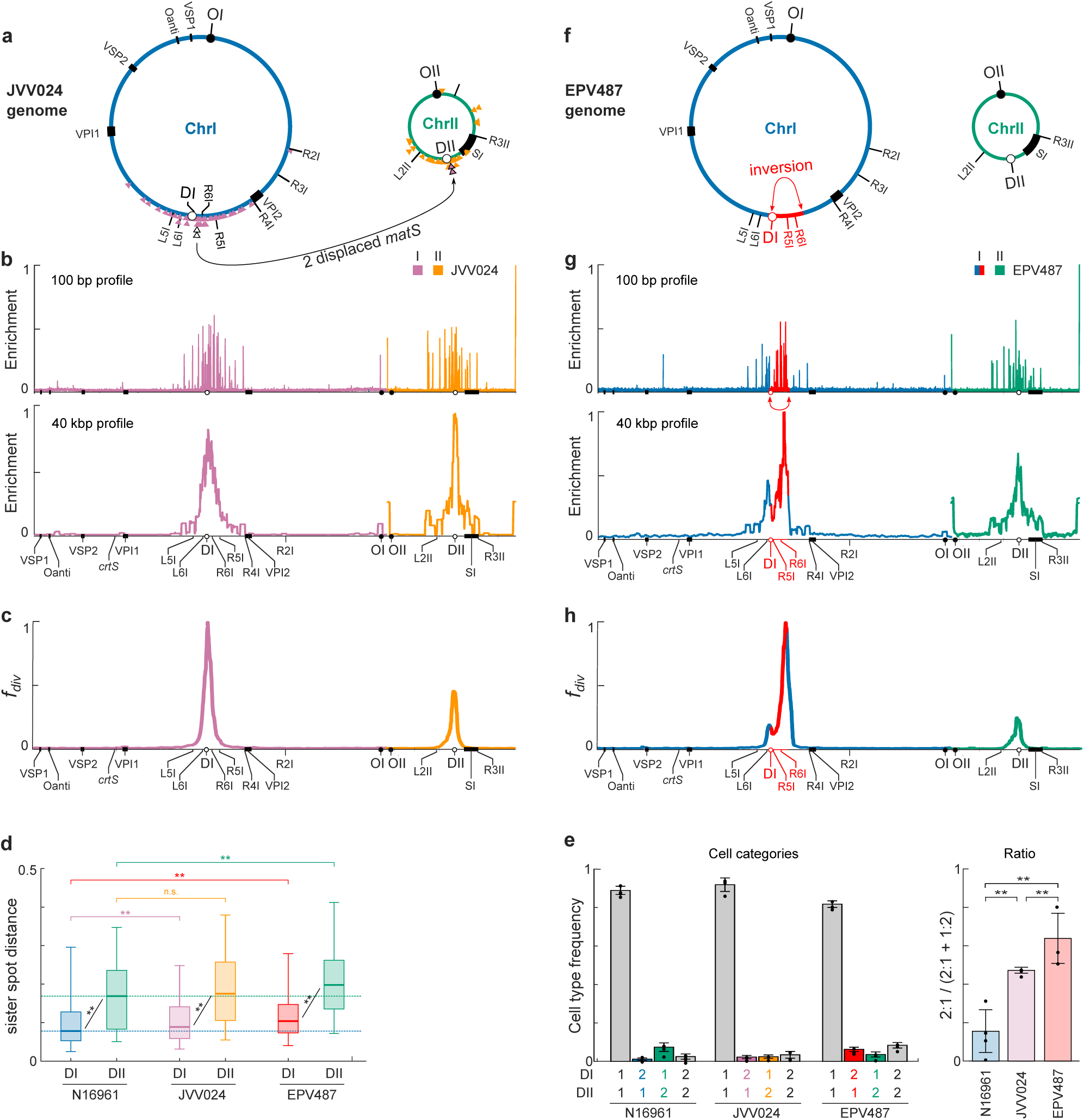
MatP action depends on the number and local density of *matS* sites. **a** and **f** JVV024 and EPV487 chromosome maps, respectively. ChrI and ChrII are depicted in blue and green, respectively. The position of the *matS* sites on JVV024 ChrI and ChrII are indicated by pink and orange triangles, respectively. The two *matS* sites that were deleted from JVV024 ChrI and inserted in ChrII are depicted by white and pink triangles with a black contour, respectively. The inverted region in EPV487 *TerI* is highlighted in red. Genomic features of interest are indicated as in Fig. 1b. **b** and **g** ChIP-seq analysis of the binding of *V. cholerae* MatP in JVV024 and EPV487, respectively. The 100 bp and 40 kbp sliding average profiles are shown to indicate the *matS* sites and to highlight the enrichment of MatP in *TerI* and *TerII*. Genomic features of interest are displayed on the x-axis. **c** and **h** FtsK/XerCD/*dif1*-based Hi-SC2 analysis of the relative frequency of contacts of sister copies during septum closure in JVV024 and EPV487, respectively. Results are shown at a 40 kbp resolution. Genomic features of interest are displayed on the x-axis. **d** Distance along the longitudinal axis between sister copies of DI and DII spots. The median (horizontal bar), the 25th and the 75th percentiles (open box) and the 5th and the 95th percentiles (error bars) of the sister spot distances are represented for each sample. Results correspond to 4 (N16961) and 3 (JJV024 and EPV487) biological replicates, with a median number of 254 cells per strain per replicate (Supplementary Data 3). **\*\*** and n.s.: significantly and not significantly different, respectively (Wilcoxon-Mann-Whitney test, Supplementary Table 1). **e** Cells were classified in four categories as a function of the number of DI and DII spots (1 DI & 1 DII spots, 2 DI &1 DII spots, 1 DI & 2 DII spots, and 2 DI & 2 DII spots) and the ratio of cells from the second category over the number of cells from the second and third categories (mean and SD). Results correspond to 4 (N16961) and 3 (JJV024 and EPV487) biological replicates, with a median number of 3031 cells per strain per replicate (Supplementary Data 3). **\*\*** and n.s.: significantly and not significantly different, respectively (Wilcoxon-Mann-Whitney test, Supplementary Table 2). Source data of panel d and e are provided as a Source Data file.

### MatP acts independently of cell division

A plasmid with two *matS* sites accumulates at mid-cell in a MatP dependent manner in *E. coli*^15^. The localisation pattern of the plasmid was shown to be due to the formation of a proteinaceous bridge between the tetramerization domain of MatP, ZapB, ZapA and FtsZ ^15,18^. In *V. cholerae*, DI localised at mid-cell at the time of cell division, consistent with the idea that MatP coordinated chromosome segregation and cell division by tethering MatP domains to the divisome (Fig. 3a and Supplementary Fig. 5a). However, DII did not localise at mid-cell at the time of division (Fig. 3a and Supplementary Fig. 5a) even though it is flanked by 34 *matS* sites (Fig. 1c). Therefore, we decided to explore the role of the tetramerization domain of MatP, ZapB and ZapA in the positioning of DI and DII.

To this end, we constructed an allele of *matP* truncated from the extreme C-terminus, which based on the homology with *E. coli matP* should prevent the formation of MatP tetramers, *matP* Δ*tetra*. MatP Δtetra had the same binding profile as WT MatP (Supplementary Fig. 8a). We then compared the cellular position of DI and DII in parallel with that of OI in WT, Δ*matP*, *matP* Δ*dimer*, *matP* Δ*tetra*, Δ*zapA* and Δ*zapB* N16961 strains in snapshot images. Demographs suggested that the choreography of segregation of DI and DII in *matP* Δ*tetra*, Δ*zapA* and Δ*zapB* cells was similar to that of WT cells, in contrast to that of Δ*matP* and *matP* Δ*dimer* cells (Supplementary Fig. 8b). To quantify the effect of the *matP* Δ*tetra*, Δ*zapA* and Δ*zapB* mutations on the cellular position of DI and DII, we analysed the distance separating sister DI copies and sister DII copies. Whereas the distance between sister DI copies was as elevated as that of sister DII copies in Δ*matP* and *matP* Δ*dimer* cells, it was significantly lower than that of DII in the *matP* Δ*tetra*, Δ*zapA* and Δ*zapB* mutants (Fig. 6a, Supplementary Table 3). However, we noticed that the distance between sister DI copies was slightly – yet significantly - elevated in the *matP* Δ*tetra*, Δ*zapA* and Δ*zapB* (Fig. 6a, Supplementary Table 3). Hence, we decided to follow the choreography of segregation of DI by time-lapse video microscopy. Results recapitulated those of the demographs. In the presence of MatP, DI localised at the new pole of newborn cells and migrated to mid-cell at the very end of the cell cycle, with sister DI copies remaining associated until the very end of the cell cycle (Fig. 6b). In Δ*matP* cells, DI was randomly distributed between the new pole and mid-cell at birth and the ¼-¾ part of the cell at the end of the cell cycle (Fig. 6b). Inspection of individual traces further showed that sister DI copies separated prior to cell division with a distance of over a quarter of the cell length (Supplementary Fig. 9b). Cells carrying the *matP* Δ*dimer* allele had the same phenotype as Δ*matP* cells (Fig. 6b). In *matP* Δ*tetra*, Δ*zapA* and Δ*zapB* cells, the fluorescence signal was less marked at the new pole of new born cells than in WT cells (Fig. 6b). Nevertheless, the choreography of segregation remained similar to that of WT cells (Fig. 6b).

**Fig. 6.**
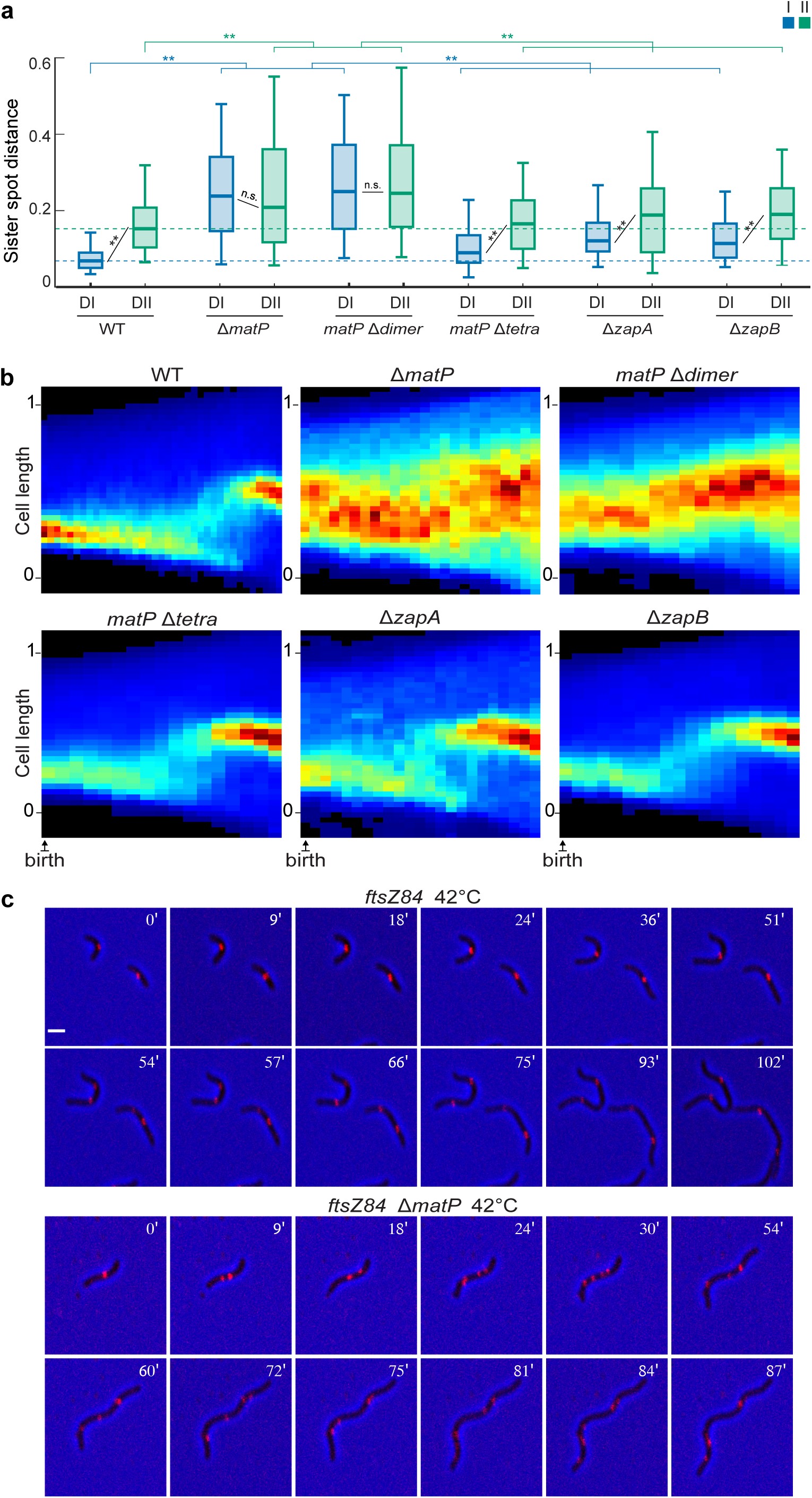
MatP acts independently of cell division and tetramer formation. **a** Relative distance between sister copies of the DI (blue) and DII (green) locus. The median (horizontal bar), the 25th and the 75th percentiles (open box) and the 5th and the 95th percentiles (error bars) of the sister spot distances are represented for each sample. Dotted lines highlight the median DI (blue) and DII (green) sister spot distance in WT cells. A median number of 225 cells were analysed for qeach strain (Supplementary data 3). **\*\*** and n.s.: significantly and not significantly different, respectively (Wilcoxon-Mann-Whitney test, Supplementary Table 3). Source data are provided as a Source Data file. **b** Cellular localisation of DI foci over-time in time-lapse experiments of WT and mutant *V. cholerae* strains. Each panel shows the compilation of individual traces. In the heat maps, black corresponds to the lowest and dark red to the highest intensities. y-axis: position along the cell length, with 0 corresponding to the new pole and 1 to the old pole. x-axis: relative advancement of the cell cycle from the first image observed after birth and the last image before division. **c** Cellular localisation of DI foci over-time in time-lapse experiments in *ftsZ84*^ts^ and *ftsZ84*^ts^ Δ*matP* cells grown at 42°C. On the top-right corner of each frame is indicated the time in minutes from the beginning of imaging. Scale bar = 2 μm.

To rule out the possibility that another cell division protein was involved in the action of MatP on the positioning of *TerI* and *TerII* at mid-cell, we compared the choreography of DI segregation in *matP ^+^* and Δ*matP* cells carrying a thermosensitive allele of *ftsZ*, *ftsZ84*^ts^. Growth of *ftsZ84*^ts^ strains at the non-permissive temperature (42°C) prevents the formation of the FtsZ ring, which leads to the dispersal of the other cell division proteins in the cytoplasm of the cell and filamentation^5^. The binding profile of MatP on the genome of *ftsZ84*^ts^ cells grown at 42°C was identical to the binding profile of MatP on the genome of WT N16961 cells and *ftsZ84*^ts^ cells grown at the permissive temperature (30°C) (Supplementary Fig. 9c). However, DI copies separated from each other at a much later stage in the *ftsZ84*^ts^ strain grown at 42°C than in its Δ*matP* derivative (Fig. 6c, Supplementary Movies 1 and 2).

Taken together, these results showed that MatP was able to keep sister *dif1* sites together at mid-cell independently of tetramer formation and divisome anchoring.

## Discussion

### *TerI* and *TerII* both contain a high density of *matS* sites

The *V. cholerae matS* sites we identified are similar to the *E. coli matS* sites, but more diverse (Fig 1, Supplementary Fig. 1), which explains why only 34 and 14 *matS* sites were predicted on ChrI and ChrII based on the *E. coli matS* motif^13^. The relative height of the ChIP-seq profile at the location of the *matS* sites was similar in the different ChIP-seq experiments we performed in the course of this study (Fig. 1, 4, 5, Supplementary Fig. 8 and 9). The ChIP-seq profile tended to be higher at the centre of the *TerI* and *TerII* MDs. This was not linked to any bias in the identity score of the sites at the centre of the domains (Fig. 1, Supplementary Data 1). The MCH1 and EPV487 data further ruled out that it was linked to the timing of replication and the action of FtsK (Fig. 4 and 5). Instead, it seemed linked to the increase in the local density of *matS* sites at the centre of the *TerI* and *TerII* MDs (Fig. 1). The presence of a *matS* site in a phage satellite that is integrated in two copies next to *dif1* (TLCΦ), in several cassettes of the super-integron, which is located next to *dif2*, and in the coding sequence of genes, suggests that there is a selection pressure to maintain the number and local density of *matS* sites. We detected additional *matS* binding sites on the genome of *V. cholerae* that were not enriched in the ChIP-seq experiments despite a good identity score (Supplementary Fig. 2). It suggests that cellular factors, such as other DNA binding proteins and ongoing transcription, probably hinder MatP binding at these positions.

### *TerI* and *TerII* cohesion and positioning are functionally distinct from MD formation

Our results suggest that MatP plays the same two roles *in V. cholerae* as it does in *E. coli*. On one hand, it structures *TerI* and *TerII* into macrodomains, which are isolated from the rest of the genome (Fig. 2). On the other hand, it is involved in the cohesion period of *TerI* and *TerII* sister copies behind replication forks, the persistence of cohesion until the initiation of cell division in the vicinity of *dif1* and *dif2*, and the positioning of *TerI* and *TerII* loci towards the new pole in young cells and to mid-cell in old cells (Fig. 3).

However, a clear distinction could be made on the mode of action of MatP in MD formation and segregation. The reduction of long-range *cis* contacts was as important at the border of the *TerI* and *TerII* MDs as in their central part (Fig. 2). This is consistent with *E. coli* studies that suggested that a few *matS* sites were sufficient for MatP to exclude and/or inactivate a DNA condensin, MukBEF, that favours long-range *cis* contacts by extruding ∼800 kbp intrachromosomal loops^13,14,30–33^. Future work will need to investigate the interplay between MatP and MukBEF in the structuration of the chromosomes in *V. cholerae*. In contrast, cohesion behind replication forks was lower at the border of the *TerI* and *TerII* MDs than in their central part (Fig. 3). In addition, cohesion only persisted until the initiation of cell division at the very centre of the *TerI* and *TerII* MDs (Fig. 3).

### Another factor than MatP is involved in the cohesion of *TerI* and *TerII* loci

The deletion of *matP* only partially alleviated the high post-replicative cohesion of *TerI* and *TerII* loci, suggesting the implication of another factor than MatP in the phenomenon (Fig. 3 and 4). In contrast, the deletion of *matP* almost entirely suppressed the persistence of cohesion until the initiation of cell division in the immediate vicinity of *dif1* and *dif2* (Fig. 3 and 4).

### Cohesion and positioning depend on the number and local density of *matS* sites

Loci at the centre of the *TerI* MD were the closest to the new pole in young cells and sister copies of these loci were the closest to each other and to mid-cell in old cells, in agreement with the persistence of cohesion (Fig. 3). This was not true for *TerII* loci, for which the deletion of *matP* only increased the range of the positions that they could occupy, and for which the persistence of cohesion until cell division was very limited (Fig. 3). We suspected that differences in the cohesion and cellular positioning of *TerI* and *TerII* loci were at least in part due to differences in the number and density of *matS* they contained. This hypothesis was corroborated with results in three synthetic strains in which the chromosomal distribution of the *matS* sites was altered, MCH1, JVV024 and EPV487: the Right *TerI* MD of MCH1, which harbours 26 *matS* sites, was highly cohesive whereas the Left *TerI* MD of MCH1, which harbours 12 *matS* sites, was not (Fig. 4); differences in the cohesion of sister copies of *dif1* and *dif2* loci were alleviated in JVV024, in which the number of *matS* sites in the *TerI* MD was reduced from 40 to 38 and that of the *TerII* MD was increased from 34 to 36 (Fig. 5); the frequency of sister chromatid contacts at the time of cell division followed the density of *matS* sites in MCH1 and EPV487 (Fig. 4 and 5).

### MatP-mediated positioning is independent of cell division and tetramer formation

The precise mid-cell positioning of *dif1* at the end of the replication cycle and the proximity of sister *dif1* copies after their separation were not altered in Δ*zapA* and Δ*zapB* cells (Fig. 6). Likewise, recent high throughput time-lapse observations showed that mid-cell migration of the *Ter* was barely modified in Δ*zapB E. coli* cells^17^. In the present study, we further show that MatP delays the separation of sister *dif1* copies in *ftsZ84*^ts^ cells grown at the non-permissive temperature, thereby demonstrating that mid-cell positioning is independent of any anchoring to a divisome component (Fig. 6).

MatP tetramers have also been proposed to mediate cohesion by linking *matS* sites together and/or to non-specific DNA in *E. coli*^12,16,17^. In contrast, we found that deletion of the tetramerization domain of MatP barely altered the mid-cell positioning of *dif1* and the distance of separation of sister *dif1* copies, thereby demonstrating that tetramer formation plays little or no role in the cohesion and mid-cell positioning of sister DNA copies in *V. cholerae* (Fig. 6). Instead, our results are consistent with a role for MatP in the regulation of decatenation. Due to the helical nature of double stranded DNA, (+) supercoils accumulate ahead of replication forks. The torsional stress is thought to cause the replication machinery to swivel around itself, generating right-handed interlinks that must be removed by topoisomerase IV (Topo IV). MukBEF has been shown to trail behind replication forks and recruit Topo IV in *E. coli*^13,34,35^. However, cohesion at the time of cell division was higher in the central part of the *TerI* and *TerII* MDs than at their borders (Fig. 3), in contrast to the restriction of long-range *cis* contacts (Fig. 2), suggesting that MatP could slow decatenation independently of its effect on MukBEF. The decrease in cohesion between the Right *TerI* MD and the *TerII* MD domain of MCH1 (Fig. 4) and at *dif1* in the *TerI* MD of EPV487 (Fig. 5) further suggests that the prolonged period of precatenane persistence in the central part of the *TerI* and *TerII* MDs is not related to their timing of replication nor to the oriented translocation of FtsK^36,37^, but to the high density of *matS* sites they contained. Finally, the earlier separation and greater distance between sister copies of *dif2* than sister copies of *dif1* suggest that decatenation is faster in *TerII* than in *TerI* due to a lower number and density of *matS* sites (Fig. 3). The short length of ChrII could also reduce the chances of leaving unresolved catenation links behind replication forks and lead to a tug-of-war between the ParAB2/*parS2* and MatP/*matS* systems, which could help separate sister copies of *TerII*. Consistent with this view, sister *TerII* copies in MCH1 separated later, were positioned closer to each other in old cells and to the cell pole in young cells than sister *TerII* copies in N16961 (Fig. 4).

It is likely that MatP-mediated inhibition of decatenation is also responsible for the initial positioning of sister copies of the *Ter* at mid-cell in *E. coli*. Consistent with this view, it was recently reported that sister *Ter* copies separated but remained in close proximity to mid-cell in Δ*zapA* and Δ*zapB* mutants^17^. Decatenation could be more effective in *E. coli* than in *V. cholerae* because of the lower number and density of *matS* sites in *Ter* than in *TerI* and *TerII*, which could further explain the importance of the MatP-ZapB-ZapA connection for the maintenance of sister *Ter* copies at mid-cell until the final stages of cell division.

Future work will need to develop a methodology to directly monitor the catenation links that remain to be removed at the end of the replication cycle in *E. coli* and *V. cholerae*.

### ChrII behaves as a *bona fide* bacterial chromosome from a functional point of view

ChrI is derived from the chromosome of the monochromosomal ancestor of the Vibrionales and Enterobacterales, whereas ChrII is derived from a megaplasmid^38^. From a genomic point of view, ChrII is considered as a ‘chromosome-plasmid’, or chromid, because it carries essential core genes, has a nucleotide composition close to that of ChrI, but possesses plasmid-type replication and partition systems^39^. However, we previously showed that ChrII participates to the regulation of the cell cycle because its origin proximal part is enriched in SlmA-binding sites^4^. Our ChiP-seq, 3C-seq, fluorescence microscopy and Hi-SC2 results further demonstrate that MatP structures *TerII* into a macrodomain, coordinates its migration with that of *TerI* and ensures its positioning close to mid-cell at the time of cell division, which is essential for the control of cell division by SlmA-mediated nucleoid occlusion (Fig. 1, 2, 5). In this regard, ChrII is organised and behaves as a *bona fide* bacterial chromosome. It will be interesting to check whether such a functional definition applies to other chromids.

## Methods

### Strains, plasmids and oligonucleotides

Bacterial strains, plasmids and oligonucleotides used in this study are listed in Supplementary Table 4. All strains are derivatives of an El Tor N16961 strain that was rendered competent by the insertion of *hapR* by specific transposition. They were constructed by natural transformation or conjugation with integration/excision plasmids. Strains construction is described in detail in Supplementary Information. Engineered strains were confirmed by PCR. Chromosomal loci were visualised by inserting a *parS^pMT1^* motif specifically detected by YGFP-ParB^pMT1^ or a *lacO* array specifically detected by LacI-RFPT or LacI-mCherry^40,41^. The list of the strains that were used in each figure panel are detailed in Supplementary Information.

### ChIP-seq

∼10^9^ cells from overnight cultures harbouring a functional *matP3flag* allele at the *matP* native locus were diluted into 100 mL of fresh media and were grown until exponential phase (OD_600_ 0.2). Cells were crosslinked for 30’ with formaldehyde 37% up to a final concentration of 1%. Crosslinking reactions were quenched for 15’ with Glycine 2.5 M up to a final concentration of 250 mM. Samples were centrifuged, washed with TBS (Tris-HCl pH 4.7 50 mM, NaCl 1M) and harvested by centrifugation. After, cells were lysed for 30’ at 37°C with 500 µL of lysis buffer I (sucrose 20%, Tris-HCl pH 8 10 mM, NaCl 50 mM, EDTA 10 mM and Lysozyme 1 mg/mL (w/v)). Then, 500 µL of lysis buffer II (Tris-HCl pH 4.7 50 mM, NaCl 150 mM, EDTA 1mM, Triton X100 1%, Roche Antiprotease Cocktail) was added and samples were sonicated for 30’ (Covaris, Duty cycle 5, Intensity 4, Cycle/burst 200). After sonication, samples were centrifuged at 17949 g for 30’ at 4°C. Supernatants were collected into a fresh tube, 50 µL were separated to use as control (Input). For immunoprecipitation (IP), an Anti-FLAG M2 affinity gel (Sigma) was added to the cell extracts and the volume was adjusted up to 1 mL in lysis buffer II. After an overnight incubation at 4°C on a rotating wheel, samples were centrifuged at 3824 g for 30 seconds at 4°C. Pellets were washed twice with TBS mixed with Tween 20 0.05% and three times with TBS. A 3xFLAG elution solution was prepared in buffer (Tris-HCl pH 4.7 500 mM, NaCl 1M) at a concentration of 25 µg/µL and diluted in TBS to a final concentration of 150 ng/µL. 2.5 volumes of elution solution were added per volume of IP sample and Input. Samples were incubated for 30’ at 4°C on a rotating wheel. Then, samples were eluted twice by centrifugation at 3824 g for 30 seconds at 4°C. RNase A was added to a final concentration of 1 µg/mL and samples were incubated for 1 hour at 37°C. Finally, samples were de-crosslinked by the addition of 5 µL of proteinase K (20 mg/mL) and overnight incubation at 65°C. The genomic DNA from the Input and the IP sample was extracted using a Qiagen PCR Purification Kit and Qiagen MinElute PCR Purification Kit, respectively. Illumina libraries were prepared using DNA SMART^TM^ ChIP-Seq Kit (TAKARA) following manufacture’s protocol. Purified libraries were subjected to sequencing on a NextSeq 500 (Illumina). After sequencing, barcodes and adapters were removed using Cutadapt ^42^ and files with total reads were created. Finally, the BWA software was used for the alignment of reads on genomes^43^ and the data was stored in the SAM format ^44^.

We used custom MatLab functions for plotting and analysis purposes (see Code availability). In brief, Input and ChIP read numbers were smoothed with a 101 bp sliding window, and divided by the total number of reads. Positions where the Input reads were twice higher or twice lower than the median of the 200001 Input reads surrounding them were replaced by NaN (Not a Number). We then computed the ratio of the ChIP read over the Input read at each position of each chromosome. We retrieved the 100 bp sequence surrounding genomic locations with a height 3 times higher than the median of the ChIP-seq profile. MEME analysis of the sequences using default parameters revealed the presence of a 13 bp DNA motif. We set the length of the motif to be exactly 13 bp in the MEME parameters to determine the position-specific probability matrix of the motif, and used it to scan the *V. cholerae* genome with MCAST. All of the motifs identified with MEME were retrieved in the MCAST motif table, which permitted to assign a quality score to their sequence (Supplementary Data 1). MCAST permitted to detect 11 additional sites at the positions of which the ChIP-seq profile was elevated (Supplementary Data 1, Supplementary Fig. 1d) and 66 sites with a quality score higher than the smallest quality score of Supplementary Data 1 sites at the positions of which the ChIP-seq profile was not elevated (Supplementary Data 2, Supplementary Fig. 2).

### Fluorescence microscopy

Cells were grown in M9 minimal medium supplemented with 0.2% fructose and 1 μg/mL thiamine. In this medium, cells perform a single replication/segregation cycle per division. Thus, each labelled locus gives rise to one or two spots, depending on whether it has been replicated and sister copies have separated or not ^9^. Cells were grown to exponential phase in liquid and spread on a 1% (w/v) agar pad (ultrapure agarose, Invitrogen). Snapshot images were acquired using a Prime BSI camera (Roper) attached to DM6000-B (Leica) microscope (Excitation source: HPX; Cubes with pass band 480/30 and 560/40 nm excitation filters, 505 and 600 nm dichroic mirrors, and pass band 535/40 and 635/60 nm emission filters). For time-lapse analyses, the slides were incubated at 30°C (and 42°C for *ftsZ84*^ts^ strains) and images were acquired every 3’ or 4’ using an Evolve 512 EMCCD camera (Roper Scientific) attached to an Axio Observer (Zeiss) with a CXUI spinning disk (Yokogawa) equipped with a triple band dichroic filter (Di01-405/488/561) and a pass band FF03-525/50-25 and long pass BLP01-561R-25 on a filter wheel. 488 nm and 561 nm 150 mW diode lasers (at 20% of their maximal delivery) were used for excitation. The list of the strains that were used in each figure panel are detailed in supplementary

### Snapshot image analysis

Cell outlines were generated using MicrobeTracker and spots were detected with SpotFinder^45^. Cell outlines and spot locations were manually curated. We used custom MatLab functions to analyse the cell Lists (See Code availability). In brief, cells were oriented on the basis that OI is located at the old pole of newborn cells whereas DI and DII are located at the new pole of newborn cells and that during the replication/segregation cycle, one of the two sister OI copies remains at the old pole and the other one migrates to the opposite pole, whereas sister DI and DII copies migrate to mid-cell and split just before cell division^9^. Demographic representations (demographs) were plotted to provide an indirect view of the cell cycle choreography of loci. The x- and y-axes of demographs correspond to the ranking number of cells when ordered according to their length and the longitudinal coordinate of the cells, respectively. Taking into account that cell length increases exponentially during the cell cycle, the ranking number can be used to estimate the advancement of the cell cycle ^5^. The numbers of cells analysed for each demograph is indicated in Supplementary Data 3.

The ranking number was used to determine the length of cells that had reached 33% and 75% of the cell cycle for each of the analysed strains. The young cells of each tagged locus were defined as the cells with a length smaller than the 33% cell cycle length in which the fluorescence label of the focus gave rise to a single spot, and the old cells as those with a length longer than the 75% cell cycle length in which the fluorescence label of the focus gave rise to two spots. The number of young and old cells for each locus of the analysed strains is indicated in Supplementary Data 3.

### Time-Lapse analysis

To follow the positions of labelled loci in the course of the cell cycle, we performed time lapse observations of cells growing in microcolonies. Cell contours were detected and cell genealogies were retraced with a MatLab script based on MicrobeTracker ^4^. After the first division event, the new pole and old pole of cells can be unambiguously attributed based on the previous division event. We used a custom MatLab function to create heat map representations of the position of labelled loci (See Code availability). In brief, the heat maps were built using cells that were born and divided in the course of the time lapse observations, i.e cells that had completed their cell cycle. The number of such cells is indicated in Supplementary Fig. 9a for each strain. Note that, because of the time interval between time points, the first image of the time lapse of these cells does not correspond to their exact timing of birth and their last image does not correspond to the exact timing of its division. The single traces were aligned using the first image following birth and the last image before division. Fluo projections were then rescaled to accommodate for differences in the length of the cells in the first image and last image of their cell cycle and in the time interval between the first image and the last image of their cell cycle. The x-axis corresponds to the relative advancement of the cell cycle from the first image after birth to the last image after birth.

### Hi-SC2

Hi-SC2 uses paired-end sequencing to determine the status and position of site-specific recombination cassettes that have been randomly-inserted by transposition^46^. The cassettes are composed of two directly repeated recombination sites separated by a DNA segment that is too short to allow their excision by intramolecular recombination. However, intermolecular recombination events between sister copies of a cassette can lead to the formation of a copy with a single recombination site. Therefore, the excision frequency of the cassette, *f*, can be used as a proxy of the length of time during which sister copies remain in contact. The local frequency of sister chromatid contacts behind replication forks can be monitored using cassettes based on the Cre/*loxP* site-specific recombination system, whose activity is independent of the cell cycle^47,48^. The local frequency of sister chromatid contacts at the onset of constriction can be monitored using cassettes based on the XerCD/*dif1* site-specific recombination system, whose activity depends on a direct contact with a cell division protein, FtsK^8,10^. Hi-SC2 transposon libraries were prepared ^49^. Briefly, transposons harbouring *loxP* or *dif1* cassettes and a kanamycin resistance gene were introduced in strains conditionally expressing Cre or XerC, respectively. Tight control of the recombinase production was achieved by placing their genes under an arabinose promoter and an antisense β-galactosidase promoter. After overnight growth on LB plates containing kanamycin and IPTG, ∼400.000 colonies were scraped, mixed in LB with IPTG and stored in aliquots at -80°C. Aliquots of the transposon library were thawed on ice and ∼10^9^ cells were diluted into 100 mL of media supplemented with IPTG. Cells were grown until exponential phase (OD_600_ 0.2). Then, ∼10^9^ cells were centrifuged and transferred into 10 mL of fresh media without IPTG. After 30’, arabinose was added at a final concentration of 0.02%. ∼10^9^ cells were recovered and frozen in liquid nitrogen after 5 hours of induction for *dif1* cassettes and 90’ for *loxP* cassettes. Genomic DNA was subjected to pair-end sequencing on a NextSeq 500 (Illumina) to determine the position of the reporters and their recombination status. Sam files were generated using Cutadapt and BWA^42–44^. The data were analysed with custom MatLab functions (see Code availability).

### Generation of 3C libraries

3C libraries were generated as described^24^. In brief, cultures were crosslinked with formaldehyde (3%). Fixed cells were lysed with SDS (0.5%) in the presence of lysozyme (35 U/mL; Tebu Bio). Their genome was subjected to HpaII digestion and re-ligation. The DNA was then purified and quantified on gel using QuantityOne software (BioRad). The resulting 3C library was sheared using a Covaris S220 instrument (Duty cycle 5, Intensity 5, cycles/burst 200, time 60 s for 4 cycles) and sheared DNA fragments were ligated to custom-made MM76 and MM77 adapters^50^. Paired-end sequencing was performed on an Illumina sequencer according to the manufacturer instructions (PairedEnd DNA sample Prep Kit – Illumina – PE-930-1001). The custom-made adapters were removed from the reads using Cutadapt. Data were then analysed using HiC-Pro^51^. Visualisation of contact-maps and computation of their ratio was performed as described in^14^. We used custom MatLab functions to plot the 3C-seq matrices (see Code availability).

## DATA AVAILABILITY

The NGS plasmid library verification, integration and excision data generated in this study have been deposited in the NIH’s Sequence Read Archive (SRA) database under accession code GSM8423721, GSM8423722, GSM8423723, GSM8423724, GSM8423725, GSM8423726, GSM8423727, GSM8423728, GSM8423729, GSM8423730, GSM8423731, GSM8423732, GSM8423733, GSM8423734, GSM8423735, GSM8423736, GSM8423737, GSM8423738, GSM8423739, GSM8423740, GSM8423741, GSM8423742, GSM8423743, GSE273190 GSM8423707, GSM8423708, GSM8423709, GSM8423710, GSM8423711, GSM8423712, GSM8423713, GSM8423714, GSM8423715, GSM8423716, GSM8423717, GSM8423718, GSM8423719, GSM8423720, GSE273189, GSM8423705, GSM8423706, GSE273188. For microscopy images, representative datasets of raw images and all metadata generated in this study have been deposited in the Mendeley public repository (https://data.mendeley.com/datasets/782fzwyc2d/1). Considering the volume of imaging data generated in this study, the remaining raw images are available from the corresponding author within two weeks upon request. The data presented in Figure 3a-d, 4c-d, 5ed, 6a, Supplementary Figure 5a and 6d are provided in the Supplementary Information/Source Data file.

## CODE AVAILABILITY

Custom MatLab functions used for analysing and plotting ChIP-seq, 3C-seq, Hi-SC2 and fluorescence microscopy results along with a set of example files are deposited in the Zenodo public repository (https://zenodo.org, DOI 10.5281/zenodo.13236102).

## ACKNOWLEDGEMENTS

The work was supported by the Agence Nationale pour la Recherche [2018-CE12-0012-03 and 2023-CE12-0018-01 to F.-X.B., 2019-CE35-0013-01 to E.G.] and the Fondation pour la Recherche Médicale [EQU202003010328 to F.-X.B.]. We thank E. Paly for helping in constructing strains and plasmids, J. Provan, M. Seba, V. Lioy, S. Duigou, and Y. Yamaichi for technical advice and helpful discussions.

## AUTHOR CONTRIBUTIONS

J.-M. D. and C. P. performed the ChIP-seq experiments. F.-X. B. analysed ChIP-seq results. M.-E. V., D. M., M. M. and R. K. performed and analysed the 3C-seq experiments. E. G. performed and analysed the fluorescence microscopy observations. E.E. and J.C. performed and analysed the Hi-SC2 experiments. E. G. and F.-X. B. supervised the work. All the authors contributed to the writing.

## COMPETING INTERESTS

The authors declare no conflict of interest.

## Notes

### Competing Interest Statement

The authors have declared no competing interest.

### Summary of Updates

Methods and figure 1, figure 2, figure 3, figure 4 and figure 5 and 6 have been clarified. Additional data has been added in supplementary figures. The text has been amended accordingly.

