## Supplementary information for "MatP local enrichment delays segregation independently of tetramer formation and septal anchoring in *Vibrio cholerae*"

### SUPPLEMENTARY FIGURES

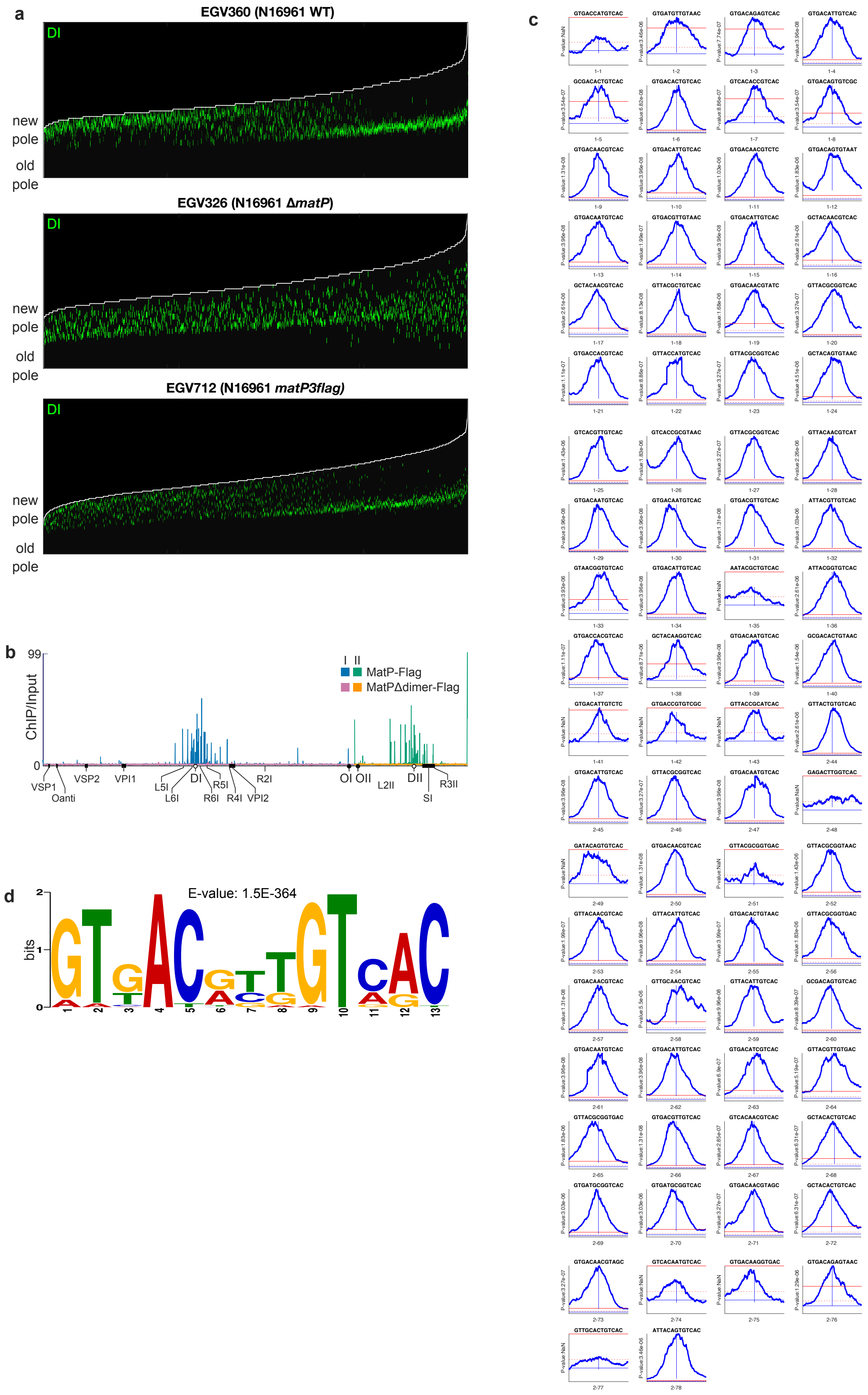

**Supplementary Fig. 1 | Numerous *matS* sites flank both *dif1* and *dif2*.**

**a** Demograhs showing the cellular location of DI in WT, Δ*matP* and *matP3flag* N16961 cells. Compilation of 2583, 2033 and 2846 cells (Supplementary Table 3). **b** Comparison of the ChIP-seq results obtained with the *matP* Δ*dimer* allele with a C-terminal triple FLAG fusion (Δ*dimer3flag*) and the *matP3flag* allele. The profiles shown are the ratio of the number of reads of ChIP and Input sequences at each chromosomal location. Read counts were smoothed using a sliding average distance of 100 bp. **c** Consensus sequence of *V. cholerae* MatP binding, built using the Meme suite based on the ChIP-seq sequences. **d** MatP3Flag ChIP-seq profile around each individual identified *matS* site listed in Supplementary Table 1. The dotted and plain red lines indicate the median of the ChIP-seq profile and the threshold limit that was used to search for MatP binding *matS* sites. The motif sequence, position order from right to left on the indicated replicon (1: ChrI; 2: ChrII), and MEME p-value are indicated. NaN: sites identified with MCAST.

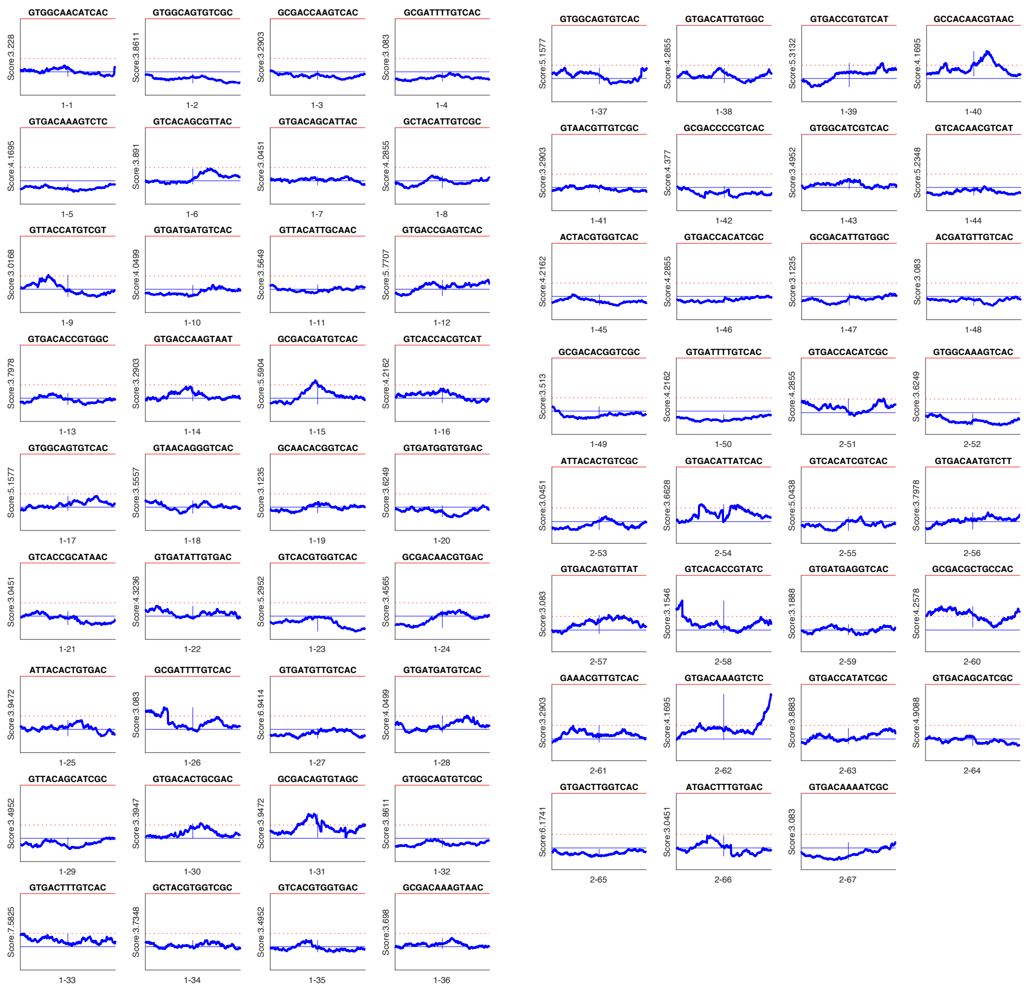

**Supplementary Fig. 2 | N16961 *V. cholerae matS* sites not bound by MatP.**

MatP3Flag ChIP-seq profile around each individual motif listed in Supplementary Table 2. The dotted and plain red lines indicate the median of the ChIP-seq profile and the threshold limit that was used to search for *matS* sites. The motif sequence, position order from right to left on the indicated replicon (1: ChrI; 2: ChrII), and MCAST score are indicated.

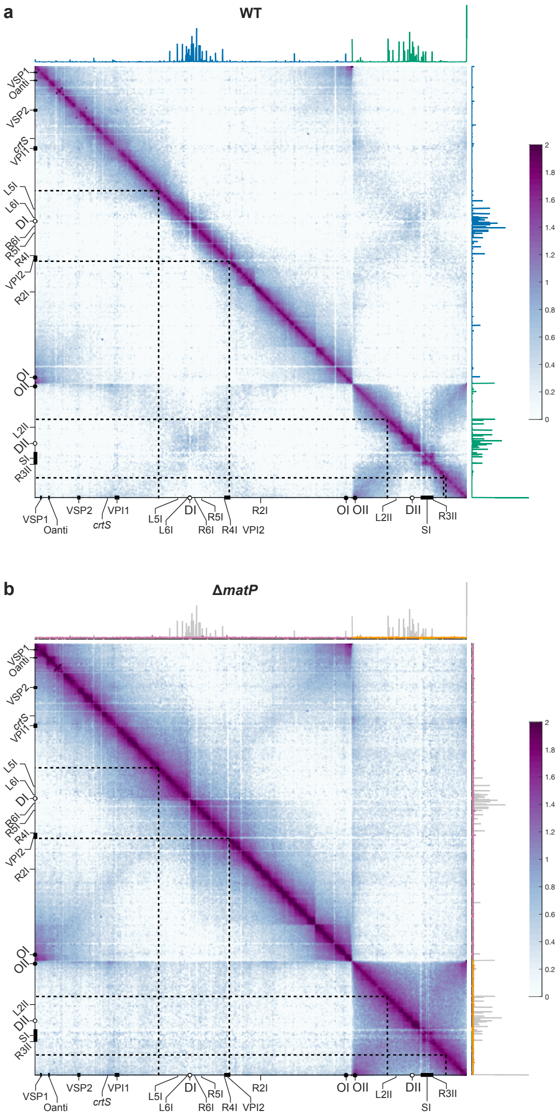


**Supplementary Fig. 3 | MatP structures *TerI* and *TerII* into macrodomains.**

Normalized contact map obtained from asynchronous populations (minimal medium, 5 kbp resolution) of WT **a** and Δ*matP* **b** *V. cholerae* cells**.** The colour scale reflects the frequency of contacts between two regions of the genome, from white (rare contacts) to dark purple (frequent contacts). Genomic features of interest are indicated on the bottom and left axes. The ChIP-seq profile (blue: ChrI; green: ChrII) obtained with the *matP3flag* allele is shown on the top and right axes of the WT 3C-seq map. The ChIP-seq profiles obtained with the *matP3flag* allele (grey) and the *matP* Δ*dimer3flag* allele (pink: ChrI, orange: ChrII) are shown on the top and right axes of the Δ*matP* 3C-seq map. Black dotted lines indicate the boundaries of the *TerI* and *TerII* MDs.

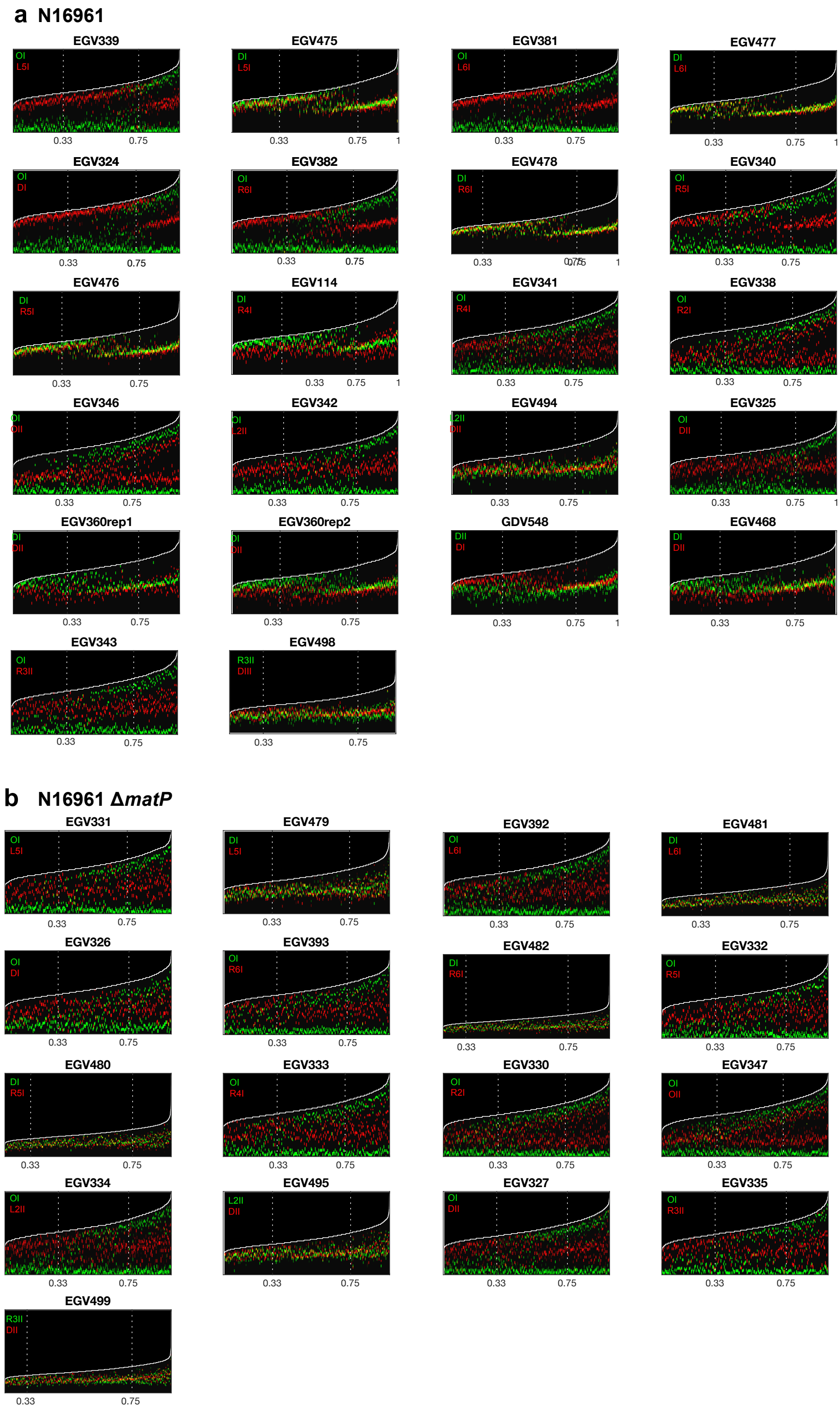


**Supplementary Fig. 4| MatP has a minor role in the positioning of ChrII loci.**

Demographs showing the cellular location of the tagged loci in individual WT **a** and Δ*matP* **b** N16961 strains. A median number of 2426 cells were used to create the demographs (Supplementary Table 3).

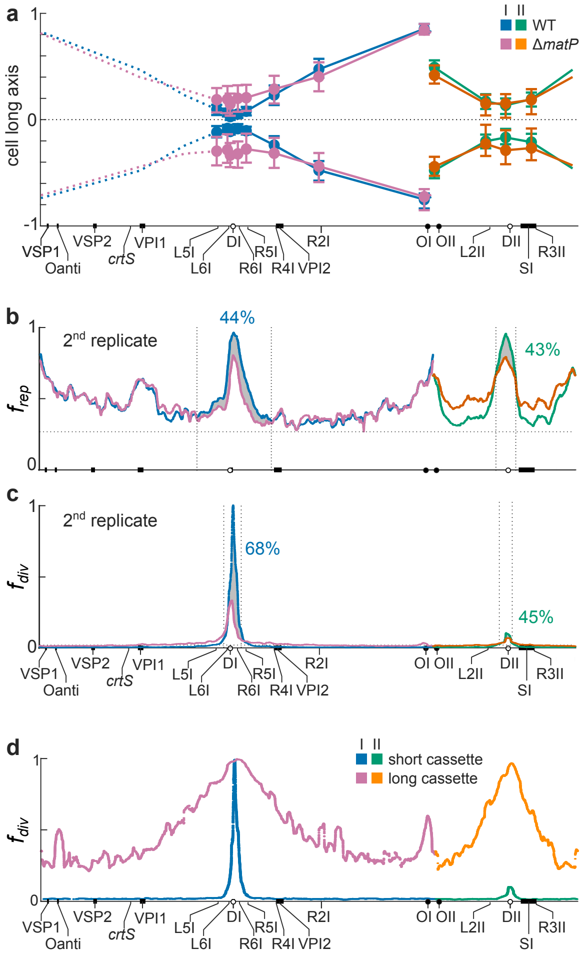


**Supplementary Fig. 5 | MatP increases cohesion in the immediate vicinity of *dif1* and *dif2*.**

**a** Relative longitudinal axis position of sister spots in old cells. Results correspond to a median number of 319 cells for each locus. By convention, the longitudinal axis coordinates of the new pole and the old pole are set to 0 and 1, respectively. The median (disk), first and third quartile (horizontal marks) of the positions/sister distances are shown. Solid lines: expected position/sister distance of intermediate loci; Dashed lines: the expected symmetrical position/sister distance of loci on the left replication arm of ChrI; Blue and pink: ChrI; green and orange: ChrII; blue and green: *matP*$^{+}$ cells (WT); pink and orange: Δ*matP* cells (Δ*matP*). **b** and **c** Replicates of the Hi-SC2 results shown in Fig. 3. **d** Comparison of FtsK/XerCD-mediated recombination of short and long *dif1* cassettes, indicative of inter and intra-molecular events, respectively. Results are shown at a 40 kbp resolution.

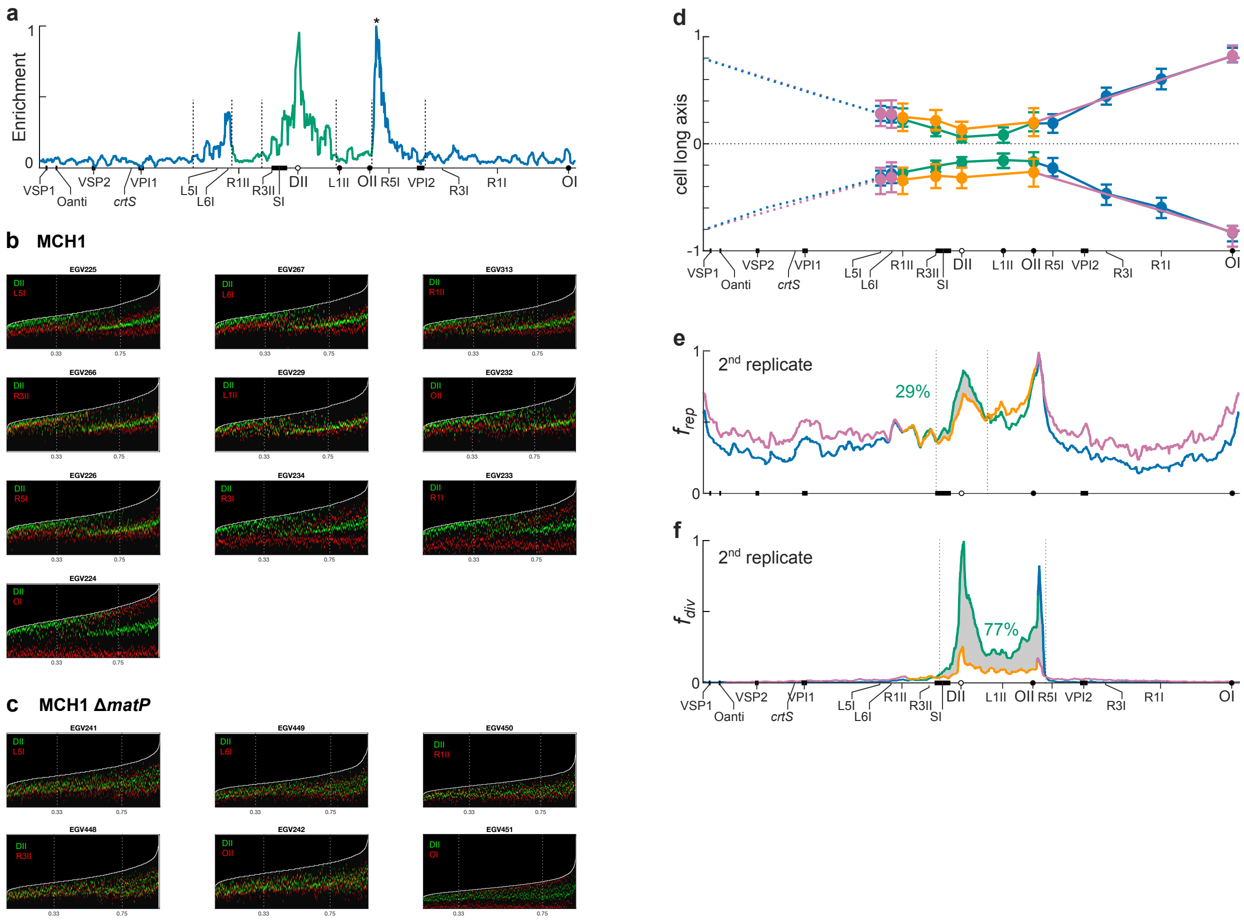


**Supplementary Fig. 6 | MatP increases cohesion in highly enriched *matS* regions.**

**a** 40 kbp sliding average of the MatP ChIP-seq data from MCH1. Blue and green lines: relative sequence enrichment profiles along the DNA originating from ChrI and ChrII, respectively. The profiles were scaled between 0 (lowest enrichment locus along the whole genome) and 1 (highest enrichment locus, indicated by a star). Genomic features of interest are displayed on the x-axis. **b** Demographs showing the cellular location of the tagged loci in individual WT MCH1 strains. A median number of 1888 cells were used to create the demographs (Supplementary Table 3). **c** Demographs showing the cellular location of the tagged loci in individual MCH1 Δ*matP* strains. A median number of 2459 cells were used to create the demographs (Supplementary Table 3). **d** Relative longitudinal axis position of each tagged locus in old cells. Results correspond to a median number of 417 cells for each locus. Cells were oriented using OI or DII. Blue and pink: regions originating from ChrI; green and orange: regions originating from ChrII. **e** and **f** Replicates of the Hi-SC2 results shown in Fig. 4.

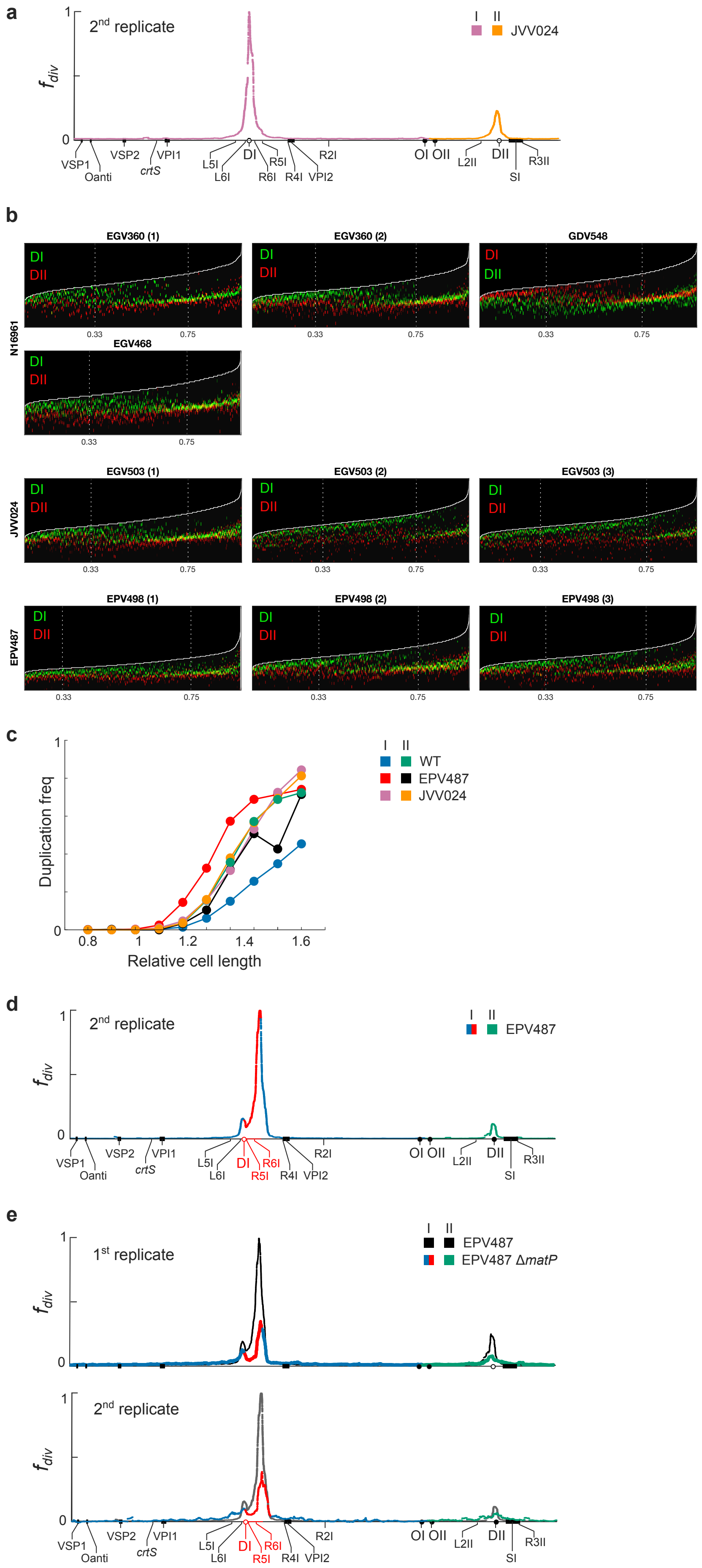

a
2nd replicate
I II
JVV024
$f_{div}$
VSP1 Oanti VSP2 VPI1 crtS L5I L6I DI R6I R5I R4I VPI2 R2I OI OII L2II DII SI R3II
b
N16961
EGV360 (1)
EGV360 (2)
GDV548
EGV468
JVV024
EGV503 (1)
EGV503 (2)
EGV503 (3)
EPV487
EPV498 (1)
EPV498 (2)
EPV498 (3)
DI
DII
0.33
0.75
c
Duplication freq
Relative cell length
I II
WT
EPV487
JVV024
d
2nd replicate
I II
EPV487
e
1st replicate
2nd replicate
I II
EPV487
EPV487 ΔmatP


70

9

**Supplementary Fig. 7 | MatP action depends on the number and local density of *matS* sites.**

**a** Replicate of the Hi-SC2 results shown in Fig. 5c. **b** Demographs showing the cellular location of DI and DII in the replicates of the fluorescence snapshot observations of N16961, JVV024 and EPV487 strains. A median number of 3031 cells were used to create the demographs (Supplementary Table 3). **c** Duplication of DI and DII loci as a function of cell length. The graph shows the frequency of cells with separated DI and DII sister copies as a function of cell length in the fluorescence snapshot observations of N16961, JVV024 and EPV487 strains. Blue, red and pink: frequency of separated DI spots in WT, EPV487 and JVV024 cells, respectively; Green, black, orange: frequency of separated DII spots in WT, EPV487 and JVV024 cells, respectively. **d** Replicate of the Hi-SC2 results shown in Fig. 5h. **e** FtsK/XerCD/*dif1*-based Hi-SC2 analysis of the relative frequency of contacts of sister copies during septum closure in EPV487 Δ*matP* cells. Results are shown at a 40 kbp resolution. ChrI and ChrII results are shown in blue and green, respectively. Results in the inverted zone of ChrI are highlighted in red. Solid black line: sister contact frequencies along the genome of EPV487 *matP*$^+$ cells from Fig. 5h.

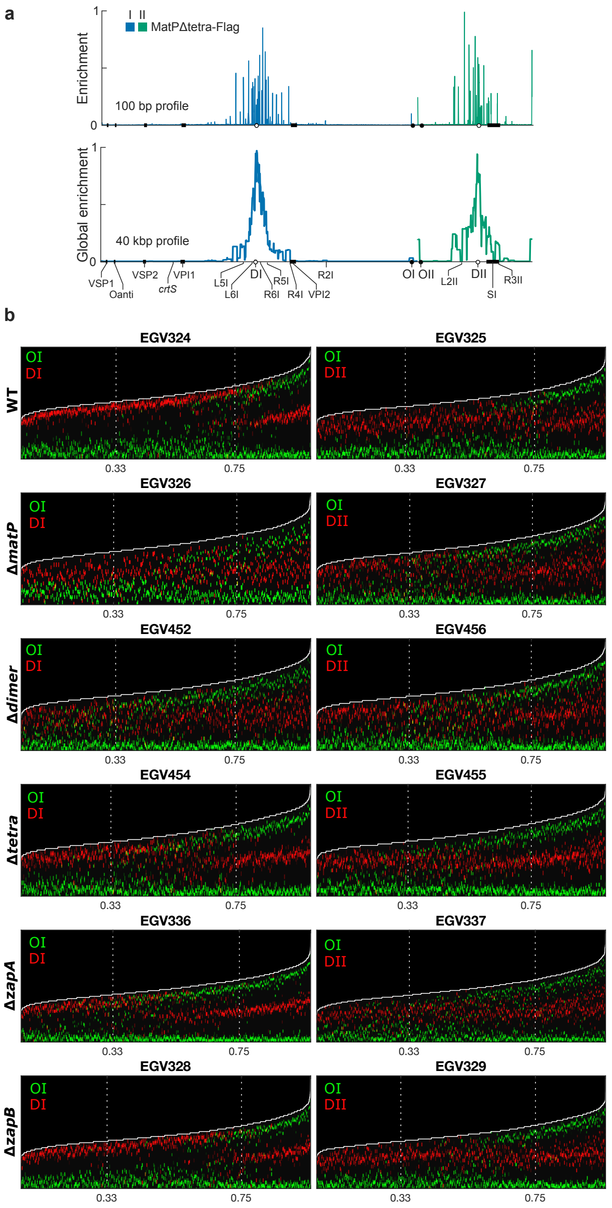


**Supplementary Fig. 8 | MatP acts independently of tetramerization.**

**a** 100 bp and 40 kbp sliding average profile of the MatP ChIP-seq data from *matP* Δ*tetra* N16961 *V. cholerae* cells. Blue/Pink and Green/Orange lines: relative sequence enrichment profiles along ChrI and ChrII, respectively. The profiles were scaled between 0 (lowest enrichment locus along the whole genome) and 1 (highest enrichment locus along the whole genome). Genomic features of interest are displayed on the x-axis. **b** Demographs showing the cellular location of OI, DI and DII in WT, Δ*matP*, *matP* Δ*dimer*, *matP* Δ*tetra*, Δ*zapA* and Δ*zapB* N16961 strains. A median number of 3174 cells were used to create the demographs (Supplementary Table 3).

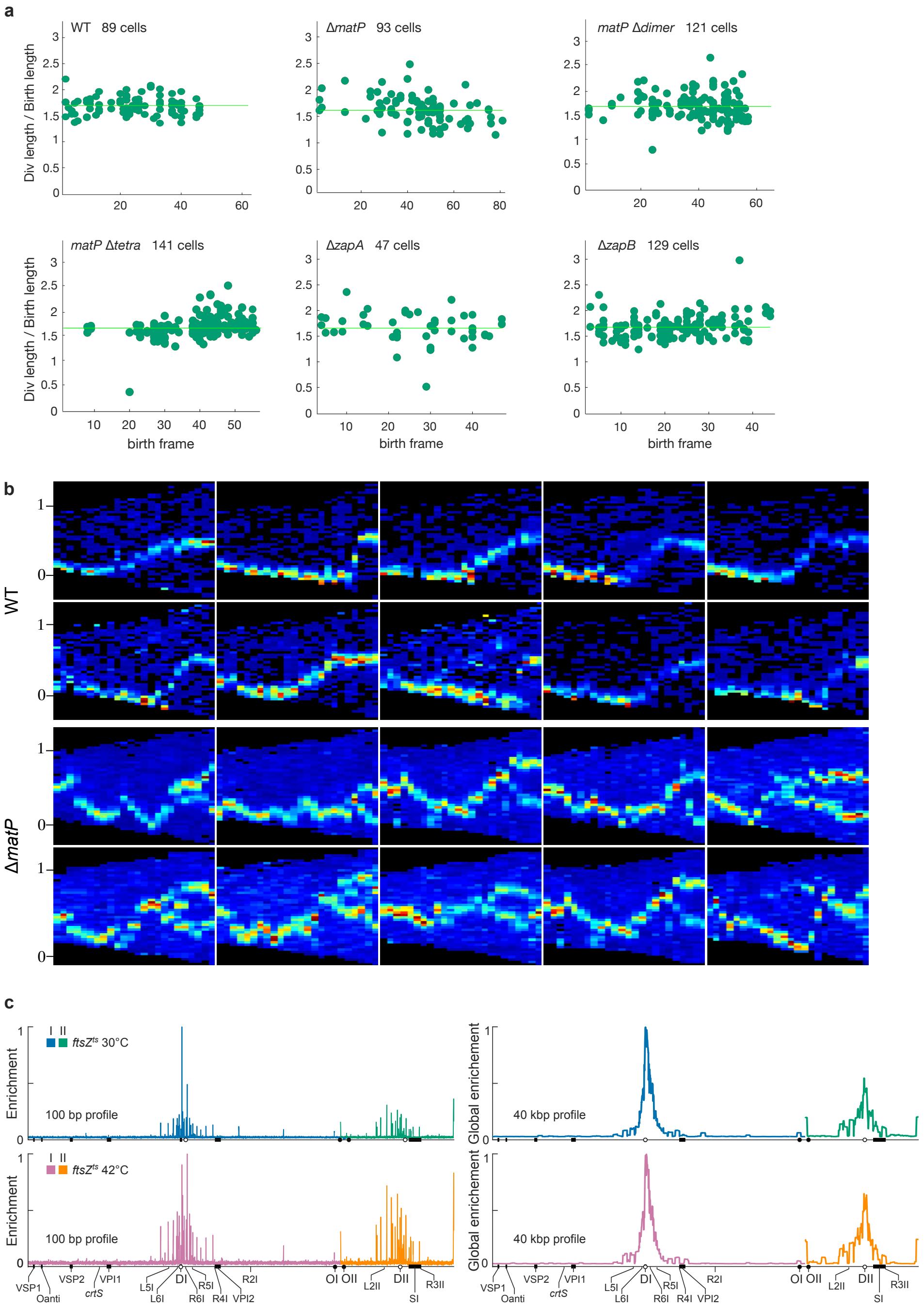


**Supplementary Fig. 9 | MatP acts independently of cell division.**

**a** In time-lapse experiments, the ratio between cell length at division (Div length) and cell length at birth (Birth length) is independent from the time point in which the cell originated (birth frame). The number of cells with complete cell cycles used to make the heat maps of Fig. 6b is indicated for each strain.

**b** Examples of the DI locus time-lapse of individual lineages in *V. cholerae* WT and Δ*matP* cells. **c** 100 bp and 40 kbp sliding average profile of the MatP ChIP-seq data from *ftsZ84*$^{ts}$

grown at 30°C (top panels) and *ftsZ84*$^{ts}$ grown at 42°C (bottom panels) N16961 cells. Blue/Pink and Green/Orange lines: relative sequence enrichment profiles along ChrI and ChrII, respectively. The profiles were scaled between 0 (lowest enrichment locus along the whole genome) and 1 (highest enrichment locus along the whole genome). Genomic features of interest are displayed on the x-axis.

### SUPPLEMENTARY TABLES

**Supplementary Table 1. Comparison of sister spot distances in N16961, JVV024 and EPV487 (Wilcoxon test P-values).** Gray shaded cells: DI sister spot distances and DII sister spot distances comparison; Blue shaded cells: DI sister spot distances comparison; Green shaded cells: DII sister spot distances comparison. P-values of significant and non-significant differences are shown in black and red, respectively.

| α = 0.001 | N16961 | JVV024 | EPV487 |
|---|---|---|---|
| N16961 | 1.1E-55 | 1.8E-05 | 1.5E-18 |
| JVV024 | 3.2E-01 | 1.3E-41 | 9.5E-7 |
| EPV487 | 6.8E-13 | 3.0E-06 | 3.4E-153 |

**Supplementary Table 2. Comparison of the ratio of number of cells with 1 DI spot and 2 DII spots vs cells with 2 DI spots and 1 DII spot in N16961, JVV024 and EPV487 (Right-tailed Wilcoxon test P-values).**

| α = 0.05 | P-value |
|---|---|
| N16961 < JVV024 | 0.0286 |
| N16961 < EPV487 | 0.0286 |
| JVV024 < EPV487 | 0.0500 |

**Supplementary Table 3. Comparison of sister spot distances in N16961, Δ*matP*, *matP* Δ*dimer*, *matP* Δ*tetra*, Δ*zapA* and Δ*zapB* (Wilcoxon test P-values).** Gray shaded cells: DI sister spot distance and DII sister spot distance comparisons; Blue shaded cells: DI sister spot distance comparisons; Green shaded cells: DII sister spot distance comparisons. P-values of significant and non-significant differences are shown in black and red, respectively.

| 0.001 | WT | dimer | matP | tetra | zapA | zapB |
|---|---|---|---|---|---|---|
| WT | 3.9E-25 | 2.4E-36 | 1.0E-35 | 4.8E-04 | 1.3E-16 | 2.0E-10 |
| dimer | 8.5E-08 | 2.0E-01 | 1.1E-01 | 1.8E-23 | 1.1E-16 | 3.0E-16 |
| matP | 6.3E-21 | 6.7E-04 | 8.2E-01 | 1.1E-23 | 4.2E-17 | 4.6E-17 |
| tetra | 4.5E-01 | 4.3E-07 | 1.7E-18 | 4.1E-10 | 8.7E-05 | 1.3E-02 |
| zapA | 2.0E-02 | 3.7E-05 | 6.6E-15 | 1.2E-01 | 4.8E-05 | 2.7E-01 |
| zapB | 7.2E-05 | 1.6E-03 | 8.7E-14 | 2.6E-03 | 2.7E-01 | 3,9E-10 |

**Supplementary Table 4**. **List of bacterial strains, plasmids and oligonucleotides**

| Strains | | |
|---|---|---|
| **Name** | **Relevant genotype or features** | **Reference** |
| B057 | N16961:: mTn7*hapR* Δ*lacZ* strep$^R$ gm$^R$ | [1] |
| C207 | N16961 *hapR*$^+$ Δ*lacZ* Δ*matP*:: *Sh ble* strep$^R$ zeo$^R$ gm$^R$ | This study |
| EEV29 | N16961 *hapR*$^+$ Δ*lacZ::*(P$_{BAD}$*::cre*-invP$_{lac}$*-Sh ble*) zeo$^R$ gm$^R$ | [2] |
| EEV92 | N16961 *hapR*$^+$ Δ*lacZ::*(P$_{BAD}$*::xerC*-invP$_{lac}$*-aad1*) Δ*matP::bla* amp$^R$ spec$^R$ gm$^R$ | This study |
| EEV94 | N16961 *hapR*$^+$ Δ*lacZ::*(P$_{BAD}$ *::cre*-invP$_{lac}$*-Sh ble*) Δ*matP::bla* zeo$^R$ gm$^R$ amp$^R$ | This study |
| EEV104 | MCH1 *hapR*$^+$ Δ*lacZ::*(P$_{BAD}$*::cre*-invP$_{lac}$*-Sh ble*) zeo$^R$ gm$^R$ | This study |
| EEV176 | MCH1 *hapR*$^+$ Δ*lacZ::*(P$_{BAD}$*::cre*-invP$_{lac}$*-Sh ble*) Δ*matP::bla* zeo$^R$ gm$^R$ amp$^R$ | This study |
| EGV27 | N16961 Δ*lacZ::*(P$_{BAD}$*::ftsZ-RFPT-Sh ble*) Δ*hapR::*(P$_{lac}$*::lacI-YGFP-cat*) + *lacO* array-*aph* inserted on ChrI at position OI zeo$^R$ gm$^R$ cml$^R$ kan$^R$ | [3] |
| EGV57 | N16961 Δ*lacZ::*(P$_{BAD}$*::ftsZ-RFPT-Sh ble*) Δ*hapR::*(P$_{lac}$*::lacI-YGFP-cat*) + *lacO* array-*aph* inserted on ChrI at position DI zeo$^R$ gm$^R$ cml$^R$ kan$^R$ | This study |
| EGV113 | N16961 *hapR*$^+$ Δ*lacZ::*(P$_{lac}$*::lacI-RFPT-YGFP-parB*$^{pMT1}$) + *parS*$^{pMT1}$ inserted on ChrI at position DI gm$^R$ | This study |
| EGV114 | N16961 *hapR*$^+$ Δ*lacZ::*(P$_{lac}$*::lacI-RFPT-YGFP-parB*$^{pMT1}$) + *lacO* array-*aph* inserted on ChrI at position R4I + *parS*$^{pMT1}$ inserted on ChrI at position DI gm$^R$ kan$^R$ | This study |
| EGV224 | MCH1 *hapR*$^+$ Δ*lacZ::*(P$_{lac}$*::lacI-RFPT-YGFP-parB*$^{pMT1}$) + *lacO* array-*aph* inserted on ChrI at position OI + *parS*$^{pMT1}$ inserted on ChrII at position DII kan$^R$ gm$^R$ | [4] |
| EGV225 | MCH1 *hapR*$^+$ Δ*lacZ::*(P$_{lac}$*::lacI-RFPT-YGFP-parB*$^{pMT1}$) + *lacO* array-*aph* inserted on ChrI at position L5I + *parS*$^{pMT1}$ inserted on ChrII at position DII kan$^R$ gm$^R$ | [4] |
| EGV226 | MCH1 *hapR*$^+$ Δ*lacZ::*(P$_{lac}$*::lacI-RFPT-YGFP-parB*$^{pMT1}$) + *lacO* array-*aph* inserted on ChrI at position R5I + *parS*$^{pMT1}$ inserted on ChrII at position DII kan$^R$ gm$^R$ | [4] |
| EGV229 | MCH1 *hapR*$^+$ Δ*lacZ::*(P$_{lac}$*::lacI-RFPT-YGFP-parB*$^{pMT1}$) + *lacO* array-*aph* inserted on ChrII at position L1II + *parS*$^{pMT1}$ inserted on ChrII at position DII kan$^R$ gm$^R$ | [4] |
| EGV232 | MCH1 *hapR*$^+$ Δ*lacZ::*(P$_{lac}$*::lacI-RFPT-YGFP-parB*$^{pMT1}$) + *lacO* array-*aph* inserted on ChrII at position OII + *parS*$^{pMT1}$ inserted on ChrII at position DII kan$^R$ gm$^R$ | [4] |
| EGV233 | MCH1 *hapR*$^+$ Δ*lacZ::*(P$_{lac}$*::lacI-RFPT-YGFP-parB*$^{pMT1}$) + *lacO* array-*aph* inserted on ChrI at position R1I + *parS*$^{pMT1}$ inserted on ChrII at position DII kan$^R$ gm$^R$ | [4] |
| EGV234 | MCH1 *hapR*$^+$ Δ*lacZ::*(P$_{lac}$*::lacI-RFPT-YGFP-parB*$^{pMT1}$) + *lacO* array-*aph* inserted on ChrI at position R3I + *parS*$^{pMT1}$ inserted on ChrII at position DII kan$^R$ gm$^R$ | [4] |
| EGV241 | MCH1 *hapR*$^+$ Δ*lacZ::*(P$_{lac}$*::lacI-RFPT-YGFP-parB*$^{pMT1}$) + *lacO* array-*aph* inserted on ChrI at position L5I + *parS*$^{pMT1}$ inserted on ChrII at position DII Δ*matP::bla* kan$^R$ gm$^R$ amp$^R$ | This study |
| EGV242 | MCH1 *hapR*$^+$ Δ*lacZ::*(P$_{lac}$*::lacI-RFPT-YGFP-parB*$^{pMT1}$) + *lacO* array-*aph* inserted on ChrII at position OII + *parS*$^{pMT1}$ inserted on ChrII at position DII Δ*matP::bla* kan$^R$ gm$^R$ amp$^R$ | This study |
| EGV266 | MCH1 *hapR*$^+$ Δ*lacZ::*(P$_{lac}$*::lacI-RFPT-YGFP-parB*$^{pMT1}$) + *lacO* array-*aph* inserted on ChrII at position R3II + *parS*$^{pMT1}$ inserted on ChrII at position DII kan$^R$ gm$^R$ | [4] |
| EGV267 | MCH1 *hapR*$^+$ Δ*lacZ::*(P$_{lac}$*::lacI-RFPT-YGFP-parB*$^{pMT1}$) + *lacO* array-*aph* inserted on ChrI at position L6I + *parS*$^{pMT1}$ inserted on ChrII at position DII kan$^R$ gm$^R$ | [4] |
| EGV313 | MCH1 *hapR*$^+$ Δ*lacZ::*(P$_{lac}$*::lacI-RFPT-YGFP-parB*$^{pMT1}$) + *lacO* array-*aph* inserted on ChrII at position R1II + *parS*$^{pMT1}$ inserted on ChrII at position DII kan$^R$ gm$^R$ | [4] |
| EGV320 | N16961 *hapR*$^+$ Δ*lacZ::*(P$_{lac}$*::lacI-RFPT-YGFP-parB*$^{pMT1}$) + *parS*$^{pMT1}$ inserted on ChrI at position OI gm$^R$ | This study |
| EGV321 | N16961 *hapR*$^+$ Δ*lacZ::*(P$_{lac}$*::lacI-RFPT-YGFP-parB*$^{pMT1}$) + *parS*$^{pMT1}$ inserted on ChrI at position OI Δ*matP::bla* gm$^R$ amp$^R$ | This study |
| EGV324 | N16961 *hapR*$^+$ Δ*lacZ::*(P$_{lac}$*::lacI-RFPT-YGFP-parB*$^{pMT1}$) + *lacO* array-*aph* inserted on ChrI at position DI + *parS*$^{pMT1}$ inserted on ChrI at position OI gm$^R$ kan$^R$ | This study |
| EGV325 | N16961 *hapR*$^+$ Δ*lacZ::*(P$_{lac}$*::lacI-RFPT-YGFP-parB*$^{pMT1}$) + *lacO* array-*aph* inserted on ChrII at position DII + *parS*$^{pMT1}$ inserted on ChrI at position OI gm$^R$ kan$^R$ | This study |
| EGV326 | N16961 *hapR*$^+$ Δ*lacZ::*(P$_{lac}$*::lacI-RFPT-YGFP-parB*$^{pMT1}$) + *lacO* array-*aph* inserted on ChrI at position DI + *parS*$^{pMT1}$ inserted on ChrI at position OI Δ*matP::bla* gm$^R$ kan$^R$ amp$^R$ | This study |
| EGV327 | N16961 *hapR*$^+$ Δ*lacZ::*(P$_{lac}$*::lacI-RFPT-YGFP-parB*$^{pMT1}$) + *lacO* array-*aph* inserted on ChrII at position DII + *parS*$^{pMT1}$ inserted on ChrI at position OI Δ*matP::bla* gm$^R$ kan$^R$ amp$^R$ | This study |
| EGV328 | N16961 *hapR*$^+$ Δ*lacZ::*(P$_{lac}$*::lacI-RFPT-YGFP-parB*$^{pMT1}$) + *lacO* array-*aph* inserted on ChrI at position DI + *parS*$^{pMT1}$ inserted on ChrI at position OI Δ*zapB::Sh ble* gm$^R$ kan$^R$ zeo$^R$ | This study |
| EGV329 | N16961 *hapR*$^+$ Δ*lacZ::*(P$_{lac}$*::lacI-RFPT-YGFP-parB*$^{pMT1}$) + *lacO* array-*aph* inserted on ChrII at position DII + *parS*$^{pMT1}$ inserted on ChrI at position OI Δ*zapB::Sh ble* gm$^R$ kan$^R$ zeo$^R$ | This study |
| EGV330 | N16961 *hapR*$^+$ Δ*lacZ::*(P$_{lac}$*::lacI-RFPT-YGFP-parB*$^{pMT1}$) + *lacO* array-*aph* inserted on ChrI at position R2I + *parS*$^{pMT1}$ inserted on ChrI at position OI Δ*matP::bla* gm$^R$ kan$^R$ amp$^R$ | This study |

| | | |
|---|---|---|
| EGV331 | N16961 *hapR*$^+$ Δ*lacZ::*($P_{lac}$*::lacI-RFPT-YGFP-parB*$^{pMT1}$) + *lacO* array-*aph* inserted on ChrI at position L5I + *parS*$^{pMT1}$ inserted on ChrI at position OI Δ*matP::bla* gm$^R$ kan$^R$ amp$^R$ | This study |
| EGV332 | N16961 *hapR*$^+$ Δ*lacZ::*($P_{lac}$*::lacI-RFPT-YGFP-parB*$^{pMT1}$) + *lacO* array-*aph* inserted on ChrI at position R5I + *parS*$^{pMT1}$ inserted on ChrI at position OI Δ*matP::bla* gm$^R$ kan$^R$ amp$^R$ | This study |
| EGV333 | N16961 *hapR*$^+$ Δ*lacZ::*($P_{lac}$*::lacI-RFPT-YGFP-parB*$^{pMT1}$) + *lacO* array-*aph* inserted on ChrI at position R4I + *parS*$^{pMT1}$ inserted on ChrI at position OI Δ*matP::bla* gm$^R$ kan$^R$ amp$^R$ | This study |
| EGV334 | N16961 *hapR*$^+$ Δ*lacZ::*($P_{lac}$*::lacI-RFPT-YGFP-parB*$^{pMT1}$) + *lacO* array-*aph* inserted on ChrII at position L2II + *parS*$^{pMT1}$ inserted on ChrI at position OI Δ*matP::bla* gm$^R$ kan$^R$ amp$^R$ | This study |
| EGV335 | N16961 *hapR*$^+$ Δ*lacZ::*($P_{lac}$*::lacI-RFPT-YGFP-parB*$^{pMT1}$) + *lacO* array-*aph* inserted on ChrII at position R3II + *parS*$^{pMT1}$ inserted on ChrI at position OI Δ*matP::bla* gm$^R$ kan$^R$ amp$^R$ | This study |
| EGV336 | N16961 *hapR*$^+$ Δ*lacZ::*($P_{lac}$*::lacI-RFPT-YGFP-parB*$^{pMT1}$) + *lacO* array-*aph* inserted on ChrI at position DI + *parS*$^{pMT1}$ inserted on ChrI at position OI Δ*zapA::arr2* gm$^R$ kan$^R$ rif$^R$ | This study |
| EGV337 | N16961 *hapR*$^+$ Δ*lacZ::*($P_{lac}$*::lacI-RFPT-YGFP-parB*$^{pMT1}$) + *lacO* array-*aph* inserted on ChrII at position DII + *parS*$^{pMT1}$ inserted on ChrI at position OI Δ*zapA::arr2* gm$^R$ kan$^R$ rif$^R$ | This study |
| EGV338 | N16961 *hapR*$^+$ Δ*lacZ::*($P_{lac}$*::lacI-RFPT-YGFP-parB*$^{pMT1}$) + *lacO* array-*aph* inserted on ChrI at position R2I + *parS*$^{pMT1}$ inserted on ChrI at position OI gm$^R$ kan$^R$ | This study |
| EGV339 | N16961 *hapR*$^+$ Δ*lacZ::*($P_{lac}$*::lacI-RFPT-YGFP-parB*$^{pMT1}$) + *lacO* array-*aph* inserted on ChrI at position L5I + *parS*$^{pMT1}$ inserted on ChrI at position OI gm$^R$ kan$^R$ | This study |
| EGV340 | N16961 *hapR*$^+$ Δ*lacZ::*($P_{lac}$*::lacI-RFPT-YGFP-parB*$^{pMT1}$) + *lacO* array-*aph* inserted on ChrI at position R5I + *parS*$^{pMT1}$ inserted on ChrI at position OI gm$^R$ kan$^R$ | This study |
| EGV341 | N16961 *hapR*$^+$ Δ*lacZ::*($P_{lac}$*::lacI-RFPT-YGFP-parB*$^{pMT1}$) + *lacO* array-*aph* inserted on ChrI at position R4I + *parS*$^{pMT1}$ inserted on ChrI at position OI gm$^R$ kan$^R$ | This study |
| EGV342 | N16961 *hapR*$^+$ Δ*lacZ::*($P_{lac}$*::lacI-RFPT-YGFP-parB*$^{pMT1}$) + *lacO* array-*aph* inserted on ChrII at position L2II + *parS*$^{pMT1}$ inserted on ChrI at position OI gm$^R$ kan$^R$ | This study |
| EGV343 | N16961 *hapR*$^+$ Δ*lacZ::*($P_{lac}$*::lacI-RFPT-YGFP-parB*$^{pMT1}$) + *lacO* array-*aph* inserted on ChrII at position R3II + *parS*$^{pMT1}$ inserted on ChrI at position OI gm$^R$ kan$^R$ | This study |
| EGV346 | N16961 *hapR*$^+$ Δ*lacZ::*($P_{lac}$*::lacI-RFPT-YGFP-parB*$^{pMT1}$) + *lacO* array-*aph* inserted on ChrII at position OII + *parS*$^{pMT1}$ inserted on ChrI at position OI gm$^R$ kan$^R$ | This study |
| EGV347 | N16961 *hapR*$^+$ Δ*lacZ::*($P_{lac}$*::lacI-RFPT-YGFP-parB*$^{pMT1}$) + *lacO* array-*aph* inserted on ChrII at position OII + *parS*$^{pMT1}$ inserted on ChrI at position OI Δ*matP::bla* gm$^R$ kan$^R$ amp$^R$ | This study |
| EGV353 | N16961 Δ*lacZ::*($P_{BAD}$*::ftsZ-RFPT-Sh ble*) Δ*hapR::*($P_{lac}$*::lacI-YGFP-cat*) + *lacO* array-*aph* inserted on ChrI at position DI Δ*matP::bla* zeo$^R$ gm$^R$ cml$^R$ kan$^R$ amp$^R$ | This study |
| EGV360 | N16961 *hapR*$^+$ Δ*lacZ::*($P_{lac}$*::lacI-RFPT-YGFP-parB*$^{pMT1}$) + *lacO* array-*aph* inserted on ChrII at position DII + *parS*$^{pMT1}$ inserted on ChrI at position DI gm$^R$ kan$^R$ | This study |
| EGV363 | N16961 *hapR*$^+$ *ftsZ84* Δ*lacZ::*($P_{lac}$*::lacI-RFPT-YGFP-parB*$^{pMT1}$*-Sh ble*) + *lacO* array-*aph* inserted on ChrI at position DI + *parS*$^{pMT1}$*-cat* inserted on ChrI at position OI gm$^R$ kan$^R$ cml$^R$ zeo$^R$ | This study |
| EGV365 | N16961 *hapR*$^+$ *ftsZ84* Δ*lacZ::*($P_{lac}$*::lacI-RFPT-YGFP-parB*$^{pMT1}$*-Sh ble*) + *lacO* array-*aph* inserted on ChrI at position DI + *parS*$^{pMT1}$*-cat* inserted on ChrI at position OI Δ*matP::bla* gm$^R$ kan$^R$ cml$^R$ zeo$^R$ amp$^R$ | This study |
| EGV381 | N16961 *hapR*$^+$ Δ*lacZ::*($P_{lac}$*::lacI-RFPT-YGFP-parB*$^{pMT1}$) + *lacO* array-*aph* inserted on ChrI at position L6I + *parS*$^{pMT1}$ inserted on ChrI at position OI gm$^R$ kan$^R$ | This study |
| EGV382 | N16961 *hapR*$^+$ Δ*lacZ::*($P_{lac}$*::lacI-RFPT-YGFP-parB*$^{pMT1}$) + *lacO* array-*aph* inserted on ChrI at position R6I + *parS*$^{pMT1}$ inserted on ChrI at position OI gm$^R$ kan$^R$ | This study |
| EGV392 | N16961 *hapR*$^+$ Δ*lacZ::*($P_{lac}$*::lacI-RFPT-YGFP-parB*$^{pMT1}$) + *lacO* array-*aph* inserted on ChrI at position L6I + *parS*$^{pMT1}$ inserted on ChrI at position OI Δ*matP::bla* gm$^R$ kan$^R$ amp$^R$ | This study |
| EGV393 | N16961 *hapR*$^+$ Δ*lacZ::*($P_{lac}$*::lacI-RFPT-YGFP-parB*$^{pMT1}$) + *lacO* array-*aph* inserted on ChrI at position R6I + *parS*$^{pMT1}$ inserted on ChrI at position OI Δ*matP::bla* gm$^R$ kan$^R$ amp$^R$ | This study |
| EGV425 | N16961 Δ*lacZ::*($P_{BAD}$*::ftsZ-RFPT-Sh ble*) Δ*hapR::*($P_{lac}$*::lacI-YGFP-cat*) + *lacO* array-*aph* inserted on ChrI at position DI Δ*zapB::aad1* zeo$^R$ gm$^R$ cml$^R$ kan$^R$ spec$^R$ | This study |
| EGV426 | N16961 Δ*lacZ::*($P_{BAD}$*::ftsZ-RFPT-Sh ble*) Δ*hapR::*($P_{lac}$*::lacI-YGFP-cat*) + *lacO* array-*aph* inserted on ChrI at position DI Δ*zapA::arr2* zeo$^R$ gm$^R$ cml$^R$ kan$^R$ rif$^R$ | This study |
| EGV448 | MCH1 *hapR*$^+$ Δ*lacZ::*($P_{lac}$*::lacI-RFPT-YGFP-parB*$^{pMT1}$) + *lacO* array-*aph* inserted on ChrII at position R3II + *parS*$^{pMT1}$ inserted on ChrII at position DII Δ*matP::bla* kan$^R$ gm$^R$ amp$^R$ | This study |
| EGV449 | MCH1 *hapR*$^+$ Δ*lacZ::*($P_{lac}$*::lacI-RFPT-YGFP-parB*$^{pMT1}$) + *lacO* array-*aph* inserted on ChrI at position L6I + *parS*$^{pMT1}$ inserted on ChrII at position DII Δ*matP::bla* kan$^R$ gm$^R$ amp$^R$ | This study |
| EGV450 | MCH1 *hapR*$^+$ Δ*lacZ::*($P_{lac}$*::lacI-RFPT-YGFP-parB*$^{pMT1}$) + *lacO* array-*aph* inserted on ChrII at position R1II + *parS*$^{pMT1}$ inserted on ChrII at position DII Δ*matP::bla* kan$^R$ gm$^R$ amp$^R$ | This study |
| EGV451 | MCH1 *hapR*$^+$ Δ*lacZ::*($P_{lac}$*::lacI-RFPT-YGFP-parB*$^{pMT1}$) + *lacO* array-*aph* inserted on ChrI at position OI + *parS*$^{pMT1}$ inserted on ChrII at position DII Δ*matP::bla* kan$^R$ gm$^R$ amp$^R$ | This study |
| EGV452 | N16961 *hapR*$^+$ Δ*lacZ::*($P_{lac}$*::lacI-RFPT-YGFP-parB*$^{pMT1}$) + *lacO* array-*aph* inserted on ChrI at position DI + *parS*$^{pMT1}$ inserted on ChrI at position OI *matP*Δdimer*::arr2* gm$^R$ kan$^R$ rif$^R$ | This study |

| | | |
|---|---|---|
| EGV454 | N16961 *hapR*$^+$ Δ*lacZ*::(P$_{lac}$::*lacI-RFPT-YGFP-parB*$^{pMT1}$) + *lacO* array-*aph* inserted on ChrI at position DI + *parS*$^{pMT1}$ inserted on ChrI at position OI *matP*Δtetra::*arr2* gm$^R$ kan$^R$ rif$^R$ | This study |
| EGV455 | N16961 *hapR*$^+$ Δ*lacZ*::(P$_{lac}$::*lacI-RFPT-YGFP-parB*$^{pMT1}$) + *lacO* array-*aph* inserted on ChrII at position DII + *parS*$^{pMT1}$ inserted on ChrI at position OI *matP*Δtetra::*arr2* gm$^R$ kan$^R$ rif$^R$ | This study |
| EGV456 | N16961 *hapR*$^+$ Δ*lacZ*::(P$_{lac}$::*lacI-RFPT-YGFP-parB*$^{pMT1}$) + *lacO* array-*aph* inserted on ChrII at position DII + *parS*$^{pMT1}$ inserted on ChrI at position OI *matP*Δdimer::*arr2* gm$^R$ kan$^R$ rif$^R$ | This study |
| EGV468 | N16961 *hapR*$^+$ Δ*lacZ*::(P$_{lac}$::*lacI-RFPT-YGFP-parB*$^{pMT1}$) + *lacO* array-*aph* inserted on ChrII at position DII + *parS*$^{pMT1}$-*cat* inserted on ChrI at position DI gm$^R$ kan$^R$ cml$^R$ | This study |
| EGV475 | N16961 *hapR*$^+$ Δ*lacZ*::(P$_{lac}$::*lacI-RFPT-YGFP-parB*$^{pMT1}$) + *lacO* array-*aph* inserted on ChrI at position L5I + *parS*$^{pMT1}$-*cat* inserted on ChrI at position DI gm$^R$ kan$^R$ cml$^R$ | This study |
| EGV476 | N16961 *hapR*$^+$ Δ*lacZ*::(P$_{lac}$::*lacI-RFPT-YGFP-parB*$^{pMT1}$) + *lacO* array-*aph* inserted on ChrI at position R5I + *parS*$^{pMT1}$-*cat* inserted on ChrI at position DI gm$^R$ kan$^R$ cml$^R$ | This study |
| EGV477 | N16961 *hapR*$^+$ Δ*lacZ*::(P$_{lac}$::*lacI-RFPT-YGFP-parB*$^{pMT1}$) + *lacO* array-*aph* inserted on ChrI at position L6I + *parS*$^{pMT1}$-*cat* inserted on ChrI at position DI gm$^R$ kan$^R$ cml$^R$ | This study |
| EGV478 | N16961 *hapR*$^+$ Δ*lacZ*::(P$_{lac}$::*lacI-RFPT-YGFP-parB*$^{pMT1}$) + *lacO* array-*aph* inserted on ChrI at position R6I + *parS*$^{pMT1}$-*cat* inserted on ChrI at position DI gm$^R$ kan$^R$ cml$^R$ | This study |
| EGV479 | N16961 *hapR*$^+$ Δ*lacZ*::(P$_{lac}$::*lacI-RFPT-YGFP-parB*$^{pMT1}$) + *lacO* array-*aph* inserted on ChrI at position L5I + *parS*$^{pMT1}$-*cat* inserted on ChrI at position DI Δ*matP*::*bla* gm$^R$ kan$^R$ cml$^R$ amp$^R$ | This study |
| EGV480 | N16961 *hapR*$^+$ Δ*lacZ*::(P$_{lac}$::*lacI-RFPT-YGFP-parB*$^{pMT1}$) + *lacO* array-*aph* inserted on ChrI at position R5I + *parS*$^{pMT1}$-*cat* inserted on ChrI at position DI Δ*matP*::*bla* gm$^R$ kan$^R$ cml$^R$ amp$^R$ | This study |
| EGV481 | N16961 *hapR*$^+$ Δ*lacZ*::(P$_{lac}$::*lacI-RFPT-YGFP-parB*$^{pMT1}$) + *lacO* array-*aph* inserted on ChrI at position L6I + *parS*$^{pMT1}$-*cat* inserted on ChrI at position DI Δ*matP*::*bla* gm$^R$ kan$^R$ cml$^R$ amp$^R$ | This study |
| EGV482 | N16961 *hapR*$^+$ Δ*lacZ*::(P$_{lac}$::*lacI-RFPT-YGFP-parB*$^{pMT1}$) + *lacO* array-*aph* inserted on ChrI at position R6I + *parS*$^{pMT1}$-*cat* inserted on ChrI at position DI Δ*matP*::*bla* gm$^R$ kan$^R$ cml$^R$ amp$^R$ | This study |
| EGV494 | N16961 *hapR*$^+$ Δ*lacZ*::(P$_{lac}$::*lacI-RFPT-YGFP-parB*$^{pMT1}$) + *lacO* array-*aph* inserted on ChrII at position DII + *parS*$^{pMT1}$-*cat* inserted on ChrII at position L2II gm$^R$ kan$^R$ cml$^R$ | This study |
| EGV495 | N16961 *hapR*$^+$ Δ*lacZ*::(P$_{lac}$::*lacI-RFPT-YGFP-parB*$^{pMT1}$) + *lacO* array-*aph* inserted on ChrII at position DII + *parS*$^{pMT1}$-*cat* inserted on ChrII at position L2II Δ*matP*::*bla* gm$^R$ kan$^R$ cml$^R$ amp$^R$ | This study |
| EGV498 | N16961 *hapR*$^+$ Δ*lacZ*::(P$_{lac}$::*lacI-RFPT-YGFP-parB*$^{pMT1}$) + *lacO* array-*aph* inserted on ChrII at position DII + *parS*$^{pMT1}$-*cat* inserted on ChrII at position R3II gm$^R$ kan$^R$ cml$^R$ | This study |
| EGV499 | N16961 *hapR*$^+$ Δ*lacZ*::(P$_{lac}$::*lacI-RFPT-YGFP-parB*$^{pMT1}$) + *lacO* array-*aph* inserted on ChrII at position DII + *parS*$^{pMT1}$-*cat* inserted on ChrII at position R3II Δ*matP*::*bla* gm$^R$ kan$^R$ cml$^R$ amp$^R$ | This study |
| EGV503 | N16961 *hapR*$^+$ Δ*lacZ*::(P$_{lac}$::*lacI-RFPT-YGFP-parB*$^{pMT1}$) + *lacO* array-*aph* inserted on ChrII at position DII + *parS*$^{pMT1}$-*cat* inserted on ChrI at position DI with 2 *matS* from ChrI moved to ChrII gm$^R$ amp$^R$ spec$^R$ kan$^R$ cml$^R$ | This study |
| EGV632 | N16961 Δ*lacZ*::(P$_{BAD}$::*ftsZ-RFPT-Sh ble*) Δ*hapR*::(P$_{lac}$::*lacI-YGFP-cat*) + *lacO* array-*aph* inserted on ChrI at position DI *matP*Δdimer::*arr2* zeo$^R$ gm$^R$ cml$^R$ kan$^R$ rif$^R$ | This study |
| EGV633 | N16961 Δ*lacZ*::(P$_{BAD}$::*ftsZ-RFPT-Sh ble*) Δ*hapR*::(P$_{lac}$::*lacI-YGFP-cat*) + *lacO* array-*aph* inserted on ChrI at position DI *matP*Δtetra::*arr2* zeo$^R$ gm$^R$ cml$^R$ kan$^R$ rif$^R$ | This study |
| EGV637 | MCH1 *hapR*$^+$ Δ*lacZ*::(P$_{BAD}$::*xerC*-invP$_{lac}$-*aad1*) spec$^R$ gm$^R$ | This study |
| EGV638 | MCH1 *hapR*$^+$ Δ*lacZ*::(P$_{BAD}$::*xerC*-invP$_{lac}$-*aad1*) Δ*matP*::*bla* amp$^R$ spec$^R$ gm$^R$ | This study |
| EGV642 | N16961 *hapR*$^+$ Δ*lacZ*::(P$_{BAD}$::*xerC*-invP$_{lac}$) with 2 *matS* from ChrI moved to ChrII spec$^R$ gm$^R$ zeo$^R$ amp$^R$ | This study |
| EGV712 | N16961 *hapR*$^+$ P$_{matP}$::*matP-3xFlag-Sh ble* Δ*lacZ*::(P$_{lac}$::*lacI-RFPT-YGFP-parB*$^{pMT1}$) + *lacO* array-*aph* inserted on ChrII at position DII + *parS*$^{pMT1}$ inserted on ChrI at position DI gm$^R$ zeo$^R$ kan$^R$ | This study |
| EPV50 | N16961 *hapR*$^+$ Δ*lacZ* gm$^R$ | [5] |
| EPV390 | N16961 ChapR Δ*lacZ*::*arr2* carrying the FtsZ84$^{ts}$ chromosomal mutation (G106S) rif$^R$ gm$^R$ | [4] |
| EPV487 | N16961 *hapR*$^+$ with inversion between 1360160 bp and *dif1* zeo$^R$ gm$^R$ | This study |
| EPV488 | N16961 *hapR*$^+$ P$_{matP}$::*matP-3xFlag-aad1* with inversion between 1360160 bp and *dif1* spec$^R$ gm$^R$ | This study |
| EPV496 | N16961 *hapR*$^+$ Δ*lacZ* P$_{matP}$::*matP-3xFlag-Sh ble* with 2 *matS* from ChrI moved to ChrII spec$^R$ gm$^R$ zeo$^R$ amp$^R$ | This study |

| | | |
|---|---|---|
| EPV498 | N16961 *hapR*$^+$ Δ*lacZ::*(P$_{lac}$*::lacI-RFPT-YGFP-parB*$^{pMT1}$*-aad1*) + *lacO* array-*aph* inserted on ChrII at position DII + *parS*$^{pMT1}$*-cat* inserted on ChrI at position DI with inversion between 1360160 bp and *dif1* gm$^R$ zeo$^R$ spec$^R$ kan$^R$ cml$^R$ | This study |
| EPV499 | N16961 *hapR*$^+$ Δ*lacZ* P$_{matP}$::*matP*Δtetra domain-*3xFlag-aad1* gm$^R$ spec$^R$ | This study |
| EPV501 | N16961 *hapR*$^+$ Δ*lacZ* P$_{matP}$::*matP*Δdimer+tetra domain-*3xFlag-aad1* gm$^R$ spec$^R$ | This study |
| GDV28 | N16961 *hapR*$^+$ Δ*lacZ::*(P$_{BAD}$*::xerC*-invP$_{lac}$*-aad1*) spec$^R$ gm$^R$ | [6] |
| GDV548 | N16961 *hapR*$^+$ Δ*lacZ::*(P$_{lac}$*::lacI-mCherry-YGFP-parB*$^{pMT1}$) + *lacO* array-*aph* inserted on ChrI at position DI + *parS*$^{pMT1}$*-cat* inserted on ChrII at position DII gm$^R$ kan$^R$ cml$^R$ | [6] |
| JMDV69 | N16961 *hapR*$^+$ Δ*lacZ* P$_{matP}$::*matP-3xFlag-Sh ble* gm$^R$ zeo$^R$ | This study |
| JMDV163 | MCH1 *hapR*$^+$ Δ*lacZ* P$_{matP}$::*matP-3xFlag-Sh ble* gm$^R$ zeo$^R$ | This study |
| JMDV182 | N16961 *hapR*$^+$ Δ*lacZ ftsZ84* P$_{matP}$::*matP-3xFlag-Sh ble* gm$^R$ zeo$^R$ rif$^R$ | This study |
| JVV013 | N16961 *hapR*$^+$ Δ*lacZ::*(P$_{BAD}$*::xerC*-invP$_{lac}$*-aad1*) with inversion between 1360160 bp and *dif1* spec$^R$ gm$^R$ | This study |
| JVV018 | N16961 *hapR*$^+$ Δ*lacZ::*(P$_{BAD}$*::xerC*-invP$_{lac}$*-aad1*) Δ*matP::bla* amp$^R$ with inversion between 1360160 bp and *dif1* spec$^R$ gm$^R$ | This study |
| JVV024 | N16961 *hapR*$^+$ Δ*lacZ* with 2 *matS* from ChrI moved to ChrII gm$^R$ spec$^R$ amp$^R$ | This study |
| MCH1 | N16961 *hapR*$^+$ fusion of ChrI and ChrII | [1] |
| **Plasmids** | | |
| **Name** | **Relevant genotype or features** | **Reference** |
| pAD20 | integration-excision vector; *sacB*; ori R6K; Tet'-*lacO* array-*aph* -'Tet cml$^R$, kan$^R$ | [5] |
| pAD39 | integration-excision vector; *sacB*; ori R6K; Tet-*parS*$^{pMT1}$- FRT-*cat*-FRT-'Tet; cml$^R$ | [5] |
| pBJ31 | integration-excision vector; *sacB*; ori R6K; P$_{BAD}$*::xerC*-invP$_{lac}$-FRT-*aad1*-FRT flanked by the upstream and downstream regions of *lacZ*; spec$^R$ cml$^R$ | [6] |
| pEE23 | P$_{BAD}$*::cre*-invP$_{lac}$-FRT-*Sh ble*-FRT flanked by the upstream and downstream regions of *lacZ*; ori pUC; amp$^R$ zeo$^R$ | [2] |
| pEG233 | P$_{BAD}$*::ftsZ-RFPT-Sh ble* flanked by the upstream and downstream regions of *lacZ*; ori pUC; amp$^R$ zeo$^R$ | [4] |
| pEG245 | P$_{lac}$*::lacI-YGFP-cat* flanked by the upstream and downstream regions of *hapR*; ori pUC; amp$^R$, cml$^R$ | [3] |
| pEG360 | *zapA*::*arr2*; ori p15a; rif$^R$ cml$^R$ | This study |
| pEG379 | *vc1489-lacO* array-*aph-vc1488*; ori pUC; cml$^R$, kan$^R$ | This study |
| pEG390 | *zapB*::*aad1*; ori pUC; spec$^R$ amp$^R$ | This study |
| pEG414 | *matP* dimerization domain::*arr2*; ori p15a; rif$^R$ cml$^R$ | This study |
| pEG416 | *lacZ-attP*$^{TLC}$*-lacZ-Sh ble* insertion in ChrI at position 1360156 bp; ori pUC; amp$^R$ zeo$^R$ | This study |
| pEG417 | Suicide vector carrying P$_{BAD}$*::xerC-xerD-xafT*; *repA*$^{ts}$; ori pSC101; cml$^R$ | This study |
| pEG418 | *matP* tetra domain::*arr2*; ori p15a; rif$^R$ cml$^R$ | This study |
| pEG419 | *matP*Δtetra domain-3xFlag-*aad1* downstream region of *matP*; ori pUC; spec$^R$ amp$^R$ | This study |
| pEG420 | *matP*Δdimer+tetra domain-3xFlag-*aad1* downstream region of *matP*; ori pUC; spec$^R$ amp$^R$ | This study |
| pEG421 | *vc1488*(containing 2 *matS* sites)-*bla* inserted between *vca0560* and *vca0561*; ori p15a; amp$^R$ cml$^R$ | This study |
| pEG423 | *vc1488*(containing 2 *matS* sites)*::aad1*; ori p15a; spec$^R$ cml$^R$ | This study |
| pEG424 | Tet-*parS*$^{pMT1}$- FRT-*cat*-FRT-Tet flanked by the upstream and downstream regions of the inverted DI; ori p15a; cml$^R$ | This study |
| pEG425 | Tet-*parS*$^{pMT1}$- FRT-*cat*-FRT-Tet flanked by the upstream and downstream regions of DI; ori p15a; cml$^R$ | This study |
| pEP70 | P$_{lac}$*::lacI-RFPT-parB*$^{pMT1}$*-YGFP*-FRT-*Sh ble*-FRT flanked by the upstream and downstream regions of *lacZ*; ori pUC; amp$^R$ zeo$^R$ | [6] |
| pEP102 | *matP-3xFlag-aad1* - downstream region of *matP*; ori R6K; spec$^R$ amp$^R$ | This study |
| pEP104 | P$_{lac}$*::lacI-RFPT-parB*$^{pMT1}$*-YGFP*-FRT-*aad1*-FRT flanked by the upstream and downstream regions of *lacZ*; ori pUC; amp$^R$ spec$^R$ | This study |
| pGD184 | *zapB*::*Sh ble*; ori pUC; zeo$^R$ amp$^R$ | This study |
| pGD243 | *matP*::*bla*; ori R6K; cml$^R$ amp$^R$ | [6] |
| pJMD18 | *matP-3xFlag-Sh ble* - downstream region of *matP*; ori R6K; zeo$^R$ amp$^R$ | This study |
| **Primers** | | |
| **Name** | **Sequence 5'-3'** | **Used for** |
| 478 | GGCAAGCTTCGGGAGATGATCAAGACTAC | DI locus |
| 503 | CCGTCTAGAGGATACGGGAGATGATCAAGACTAC | Inverted DI |
| 564 | CGCGTAGCCCAAATTTAAGCAG | DI locus |
| 595 | CATGGGTGTGCCGGAGTTTG | P$_{BAD}$*::xerC* |
| 600 | TTGCGGGTCGAACGAGGATG | *lacZ* gene |

| | | |
|---|---|---|
| 1036 | TTGCGCCATTCGATGGTGTC | $P_{BAD}$::*cre* or *xerC* |
| 1212 | GGCTACGCGCCTAACAAACG | R3II locus |
| 1213 | AACAACGCCACCACCCAACC | R3II locus |
| 1514 | TGACGAGTTCTTCTGAGCGGGACTCTGG | *lacO* array-*aph* |
| 1637 | CGGGTTGGCATGGATTGTAG | *parS*$^{pMT1}$ at DI |
| 1655 | CCCAATGCATAACGGATACC | OI locus |
| 1656 | GAGGGCGGATTATAGAGAAC | OI locus |
| 1676 | GCTCTAGAAGCGAAGAGGTGGTATAAGC | L5I locus |
| 1692 | GGACTAGTGCGCAGGATCGATTTCTAGC | R5I locus |
| 1711 | GGCGCTTTATGTCTCTGAAC | R2I locus |
| 1740 | CGACCATATTGCGTAGGTTC | OII locus |
| 1741 | GCGTGATCAGCAAATAGGTC | OII locus |
| 1750 | GCGCTACGCTGACTGAGTTC | L2II locus |
| 1751 | TGCCCAGTGTGATTATCTTG | L2II locus |
| 1764 | TGCTCGATGTCGACCATTTC | DII locus |
| 1780 | TTCTCGCTCATGGCAATTCC | R4I locus |
| 1794 | AGAATCCGCAACGTATTCCC | DI locus |
| 1812 | GGTCGTGTCCACGAACTTCC | *Sh ble* gene |
| 1839 | TGGCGTATCTACAACAAGGC | *parS*$^{pMT1}$ at DI |
| 1903 | GCTCTAGAGCGCATTCACGACATTCTAC | *zapB* gene |
| 1904 | CCGGAATTCAGTAAACGGCCTTGCATGTC | *zapB* gene |
| 1857 | TTCAACTAGTCGATGAAGCAAAGCTCGTTGTC | *matP* gene |
| 1862 | TTCAACTAGTAGGCGAGCTTCTTCAAGTAGCG | *matP* gene |
| 1948 | ATCCCAATGCAACTGCTCTC | *hapR* gene |
| 1949 | CAAAGTGCGTGATTGGACTC | *hapR* gene |
| 2001 | AGCCGGTTTATTCCGCTTTG | L6I locus |
| 2300 | CCCACTAGTTTAATTAAGTTTGTGCCCCAGTTTGC | colors at *lacZ* |
| 2804 | TTTCCGCGGGATGCGCAGAATGATCGGT | *zapA* gene |
| 2806 | TTTCAGCTGCCAATCGGCATCGCGCCTT | *zapA* gene |
| 2908 | GAATTCAGGATGCCGAAGAG | DI locus |
| 3118 | GACAGAAGCTACCCACGCAAAC | *vca0560-0561* |
| 3128 | GTCGACGGAGCTCGAATTCG | *matP3flag* |
| 3162 | GAGTATCTACTCAGCGAGGC | *matP3flag* |
| 3555 | GGATCATGTAACTCGCCTTG | *vca0560-0561* |
| 3765 | GAAGTGTTGCTTGAACATGC | ChrI *ter* inversion |
| 3771 | CTAATCTAGACGCTGTCACTGTATAACAAT | *matP3flag* |
| 3791 | AGATCTAAGCTTAGTAAAGCCCTCGCTAGACGGCATAGAGAGGATCGAATGGATA | *vc1488* gene |
| 3802 | CGACAGGTTTCCCGACTGGAAAGCGGGCAGCTCTTCAACCCGCAGCATC | *vc1488* gene |
| 3858 | GGATCCATAATACGACTCACTAT | *matPΔdimer3flag* |
| 3861 | GGTGAGATAGCGAATTCCCA | R6I locus |

### SUPPLEMENTARY METHODS

**Construction and verification of strains**

All strains are derivatives of EPV50 (El Tor N16961 rendered competent by insertion of *hapR*) or MCH1 (El Tor N16961 *hapR*$^{+}$ ChrI and ChrII fusion). All strains were constructed by integration-excision or natural transformation, a 700bp-1Kbp homology sequence upstream (UP) and downstream (DWN) of the region of interest is necessary for a double cross-over event and integration of the mutagenized DNA in the receiver strain. Both plasmids and gDNA can be used as donors in natural transformation. Genes coding for antibiotic resistance are in-between FRT sites, by conjugation with plasmid pFlp2[7] expressing a Flippase we can excise the antibiotic resistance gene from *V. cholerae* strains by recombination between FRT sites. Chromosomal loci were tagged by inserting *lacO* arrays-KanR or *parS*$^{pMT1}$-CmlR in between Tet homologies by natural transformation with pAD20 and pAD39, respectively[8]. Engineered strains were confirmed by colony PCR. The control PCR primers used for each strain are listed below. The whole genome sequence of EPV487 was confirmed by nanopore sequencing.

**EEV92**: GDV28 receiver strain; natural transformation with pGD243 for *matP::bla*, control PCR 1857+1862.

**EEV94**: EEV29 receiver strain; natural transformation with pGD243 for *matP::bla*, control PCR 1857+1862.

**EEV104**: MCH1 receiver strain; natural transformation with pEE23 for $P_{BAD}$*::cre*; control PCR 600+1036.

**EEV176:** EEV104 receiver strain; natural transformation with pGD243 for *matP::bla*, control PCR 1857+1862.

**EGV57:** EPV50 receiver strain; natural transformation with pEG233 for *ftsZ-RFPT* at *lacZ*, control PCR 528+2198; natural transformation with gDNA for *lacO* array-*aph* at DI, control PCR 1514+1794; natural transformation with pEG245 for *lacI-YGFP* at *hapR*, control PCR 1948+1949.

**EGV113:** EPV50 receiver strain; natural transformation with pEP70 for *lacI-RFPT-YGFP-parB*$^{pMT1}$ at *lacZ*, control PCR 600+2300; natural transformation with gDNA for *parS*$^{pMT1}$*-cat* at DI, control PCR 564+1794.

**EGV114:** EGV113 receiver strain; natural transformation with gDNA for *lacO* array-*aph* at R4I, control PCR 1514+1780.

**EGV241:** EGV225 receiver strain; natural transformation with pGD243 for *matP::bla*, control PCR 1857+1862.

**EGV242:** EGV232 receiver strain; natural transformation with pGD243 for *matP::bla*, control PCR 1857+1862.

**EGV320:** EPV50 receiver strain; natural transformation with pEP70 for *lacI-RFPT-YGFP-parB*$^{pMT1}$ at *lacZ*, control PCR 600+2300; natural transformation with gDNA for *parS*$^{pMT1}$*-cat* at OI, control PCR 1655+1656.

**EGV321:** EGV320 receiver strain; natural transformation with pGD243 for *matP::bla*, control PCR 1857+1862.

**EGV324:** EGV320 receiver strain; natural transformation with gDNA for *lacO* array-*aph* at DI, control PCR 1514+1794.

**EGV325:** EGV320 receiver strain; natural transformation with gDNA for *lacO* array-*aph* at DII, control PCR 1514+1764.

**EGV326:** EGV324 receiver strain; natural transformation with pGD243 for *matP::bla*, control PCR 1857+1862.

**EGV327:** EGV325 receiver strain; natural transformation with pGD243 for *matP::bla*, control PCR 1857+1862.

**EGV328:** EGV324 receiver strain; natural transformation with pGD184 for *zapB::Sh ble*, control PCR 1903+1904.

**EGV329:** EGV325 receiver strain; natural transformation with pGD184 for *zapB::Sh ble*, control PCR 1903+1904.

**EGV330:** EGV321 receiver strain; natural transformation with gDNA for *lacO* array-*aph* at R2I, control PCR 1514+1711.

**EGV331:** EGV321 receiver strain; natural transformation with gDNA for *lacO* array-*aph* at L5I, control PCR 1514+1676.

**EGV332:** EGV321 receiver strain; natural transformation with gDNA for *lacO* array-*aph* at R5I, control PCR 1514+1692.

**EGV333:** EGV321 receiver strain; natural transformation with gDNA for *lacO* array-*aph* at R4I, control PCR 1514+1780.

**EGV334:** EGV321 receiver strain; natural transformation with gDNA for *lacO* array-*aph* at L2II, control PCR 1514+1750.

**EGV335:** EGV321 receiver strain; natural transformation with gDNA for *lacO* array-*aph* at R3II, control PCR 1514+1213.

**EGV336:** EGV324 receiver strain; natural transformation with pEG360 for *zapA::arr2*, control PCR 2804+2806.

**EGV337:** EGV325 receiver strain; natural transformation with pEG360 for *zapA::arr2*, control PCR 2804+2806.

**EGV338:** EGV320 receiver strain; natural transformation with gDNA for *lacO* array-*aph* at R2I, control PCR 1514+1711.

**EGV339:** EGV320 receiver strain; natural transformation with gDNA for *lacO* array-*aph* at L5I, control PCR 1514+1676.

**EGV340:** EGV320 receiver strain; natural transformation with gDNA for *lacO* array-*aph* at R5I, control PCR 1514+1692.

**EGV341:** EGV320 receiver strain; natural transformation with gDNA for *lacO* array-*aph* at R4I, control PCR 1514+1780.

**EGV342:** EGV320 receiver strain; natural transformation with gDNA for *lacO* array-*aph* at L2II, control PCR 1514+1750.

**EGV343:** EGV320 receiver strain; natural transformation with gDNA for *lacO* array-*aph* at R3II, control PCR 1514+1213.

**EGV346:** EGV320 receiver strain; natural transformation with gDNA for *lacO* array-*aph* at OII, control PCR 1740+1741.

**EGV347:** EGV321 receiver strain; natural transformation with gDNA for *lacO* array-*aph* at OII, control PCR 1740+1741.

**EGV353:** EPV50 receiver strain; natural transformation with pEG233 for *ftsZ-RFPT* at *lacZ*, control PCR 528+2198; natural transformation with gDNA for *lacO* array-*aph* at DI, control PCR 1514+1794; natural transformation with pGD243 for *matP::bla*, control PCR 1857+1862; natural transformation with pEG245 for *lacI-YGFP* at *hapR*, control PCR 1948+1949.

**EGV360:** EGV113 receiver strain; natural transformation with gDNA for *lacO* array-*aph* at DII, control PCR 1514+1764.

**EGV363:** EPV390 receiver strain carrying the *ftsZ84*$^{ts}$ mutation, natural transformation with gDNA for *parS*$^{pMT1}$*-cat* at OI, control PCR 1655+1656; natural transformation with gDNA for *lacO* array-*aph* at DI, control PCR 1514+1794; natural transformation with pEP70 for *lacI-RFPT-YGFP-parB*$^{pMT1}$ at *lacZ*, control PCR 600+2300.

**EGV365:** EGV363 receiver strain; natural transformation with pGD243 for *matP::bla*, control PCR 1857+1862.

**EGV381:** EGV320 receiver strain; natural transformation with gDNA for *lacO* array-*aph* at L6I, control PCR 1514+2001.

**EGV382:** EGV320 receiver strain; natural transformation with pEG379 for *lacO* array-*aph* at R6I, control PCR 1514+3861.

**EGV392:** EGV381 receiver strain; natural transformation with pGD243 for *matP::bla*, control PCR 1857+1862.

**EGV393:** EGV382 receiver strain; natural transformation with pGD243 for *matP::bla*, control PCR 1857+1862.

**EGV425:** EPV50 receiver strain; natural transformation with pEG233 for *ftsZ-RFPT* at *lacZ*, control PCR 528+2198; natural transformation with gDNA for *lacO* array-*aph* at DI, control PCR 1514+1794; natural transformation with pGD184 for *zapB::Sh ble*, control PCR 1903+1904; natural transformation with pEG245 for *lacI-YGFP* at *hapR*, control PCR 1948+1949.

**EGV426:** EPV50 receiver strain; natural transformation with pEG233 for *ftsZ-RFPT* at *lacZ*, control PCR 528+2198; natural transformation with gDNA for *lacO* array-*aph* at DI, control PCR 1514+1794; natural transformation with pEG360 for *zapA::arr2*, control PCR 2804+2806; natural transformation with pEG245 for *lacI-YGFP* at *hapR*, control PCR 1948+1949.

**EGV448:** EGV266 receiver strain; natural transformation with pGD243 for *matP::bla*, control PCR 1857+1862.

**EGV449:** EGV267 receiver strain; natural transformation with pGD243 for *matP::bla*, control PCR 1857+1862.

**EGV450:** EGV313 receiver strain; natural transformation with pGD243 for *matP::bla*, control PCR 1857+1862.

**EGV451:** EGV224 receiver strain; natural transformation with pGD243 for *matP::bla*, control PCR 1857+1862.

**EGV452:** EGV324 receiver strain; natural transformation with pEG414 for *matP*Δdimer*::arr2*, control PCR 1857+1862.

**EGV454:** EGV324 receiver strain; natural transformation with pEG418 for *matP*Δtetra*::arr2*, control PCR 1857+1862.

**EGV455:** EGV325 receiver strain; natural transformation with pEG418 for *matP*Δtetra*::arr2*, control PCR 1857+1862.

**EGV456:** EGV325 receiver strain; natural transformation with pEG414 for *matP*Δdimer*::arr2*, control PCR 1857+1862.

**EGV475:** EPV50 receiver strain; natural transformation with pEP70 for *lacI-RFPT-YGFP-parB*$^{pMT1}$ at *lacZ*, control PCR 600+2300; natural transformation with gDNA for *parS*$^{pMT1}$*-cat* at DI, control PCR 1839+1637; natural transformation with gDNA for *lacO* array-*aph* at L5I, control PCR 1514+1676.

**EGV468:** EPV50 receiver strain; natural transformation with pEP70 for *lacI-RFPT-YGFP-parB*$^{pMT1}$ at *lacZ*, control PCR 600+2300; natural transformation with pEG425 for *parS*$^{pMT1}$*-cat* at DI, control PCR 1839+1637 and 2908+478; natural transformation with gDNA for *lacO* array-*aph* at DII, control PCR 1514+1764.

**EGV476:** EPV50 receiver strain; natural transformation with pEP70 for *lacI-RFPT-YGFP-parB*$^{pMT1}$ at *lacZ*, control PCR 600+2300; natural transformation with gDNA for *parS*$^{pMT1}$*-cat* at DI, control PCR 1839+1637; natural transformation with gDNA for *lacO* array-*aph* at R5I, control PCR 1514+1692.

**EGV477:** EPV50 receiver strain; natural transformation with pEP70 for *lacI-RFPT-YGFP-parB*$^{pMT1}$ at *lacZ*, control PCR 600+2300; natural transformation with gDNA for *parS*$^{pMT1}$*-cat* at DI, control PCR 1839+1637; natural transformation with gDNA for *lacO* array-*aph* at L6I, control PCR 1514+2001.

**EGV478:** EPV50 receiver strain; natural transformation with pEP70 for *lacI-RFPT-YGFP-parB*$^{pMT1}$ at *lacZ*, control PCR 600+2300; natural transformation with gDNA for *parS*$^{pMT1}$*-cat* at DI, control PCR 1839+1637; natural transformation with gDNA for *lacO* array-*aph* at R6I, control PCR 1514+3861.

**EGV479:** EGV475 receiver strain; natural transformation with pGD243 for *matP::bla*, control PCR 1857+1862.

**EGV480:** EGV476 receiver strain; natural transformation with pGD243 for *matP::bla*, control PCR 1857+1862.

**EGV481:** EGV477 receiver strain; natural transformation with pGD243 for *matP::bla*, control PCR 1857+1862.

**EGV482:** EGV478 receiver strain; natural transformation with pGD243 for *matP::bla*, control PCR 1857+1862.

**EGV494:** EPV50 receiver strain; natural transformation with pEP70 for *lacI-RFPT-YGFP-parB*$^{pMT1}$ at *lacZ*, control PCR 600+2300; natural transformation with gDNA for *parS*$^{pMT1}$*-cat* at L2II, control PCR 1750+1751; natural transformation with gDNA for *lacO* array-*aph* at DII, control PCR 1514+1764.

**EGV495:** EGV494 receiver strain; natural transformation with pGD243 for *matP::bla*, control PCR 1857+1862.

**EGV498:** EPV50 receiver strain; natural transformation with pEP70 for *lacI-RFPT-YGFP-parB*$^{pMT1}$ at *lacZ*, control PCR 600+2300; natural transformation with gDNA for *parS*$^{pMT1}$*-cat* at

R3II, control PCR 1212+1213; natural transformation with gDNA for *lacO* array-*aph* at DII, control PCR 1514+1764.

**EGV499:** EGV498 receiver strain; natural transformation with pGD243 for *matP::bla*, control PCR 1857+1862.

**EGV503:** EPV50 receiver strain; natural transformation with pEP70 for *lacI-RFPT-YGFP-*$parB^{pMT1}$ at *lacZ*, control PCR 600+2300; natural transformation with gDNA for $parS^{pMT1}$*-cat* at DI, control PCR 1839+1637; natural transformation with gDNA for *lacO* array-*aph* at DII, control PCR 1514+1764; natural transformation with pEG421 to integrate *vc1488* with its 2 *matS* sites between *vca0560* and *vca0561*, control PCR 3118+3555; natural transformation with pEG423 to delete *vc1488*, control PCR 3802+3791.

**EGV632:** EPV50 receiver strain; natural transformation with pEG233 for *ftsZ-RFPT* at *lacZ*, control PCR 528+2198; natural transformation with gDNA for *lacO* array-*aph* at DI, control PCR 1514+1794; natural transformation with pEG414 for *matP*Δdimer*::arr2*, control PCR 1857+1862; natural transformation with pEG245 for *lacI-YGFP* at *hapR*, control PCR 1948+1949.

**EGV633:** EPV50 receiver strain; natural transformation with pEG233 for *ftsZ-RFPT* at *lacZ*, control PCR 528+2198; natural transformation with gDNA for *lacO* array-*aph* at DI, control PCR 1514+1794; natural transformation with pEG418 for *matP*Δtetra*::arr2*, control PCR 1857+1862; natural transformation with pEG245 for *lacI-YGFP* at *hapR*, control PCR 1948+1949.

**EGV637:** MCH1 receiver strain; natural transformation with pBJ31 for $P_{BAD}$*::xerC*; control PCR 595+1036.

**EGV638:** EGV637 receiver strain; natural transformation with pGD243 for *matP::bla*, control PCR 1857+1862.

**EGV642:** GDV28 receiver strain; natural transformation with pEG421 to integrate *vc1488* with its 2 *matS* sites between *vca0560* and *vca0561*, control PCR 3118+3555; natural transformation with pEG423 to delete *vc1488*, control PCR 3802+3791.

**EGV712:** EGV360 receiver strain, natural transformation with pJMD18 for $P_{matP}$::*matP-3xFlag-Sh ble*, control PCR 3128+3162.

**EPV487:** EPV50 strain receiver; natural transformation with pEG416 for *lacZ-*$attP^{TLC}$*-lacZ-Sh ble* insertion in ChrI at position 1360156 bp, (blue colony), control PCR 3765+1812; conjugation with the suicide plasmid pEG417, expression with 0.1% L-Arabinose of *xerC-xerD-xafT* to induce recombination between *dif1* and $attP^{TLC}$ (white colony if inversion succeeded) and incubation at 42°C to lose the suicide plasmid[9]. The whole genome sequence of EPV487 was confirmed by nanopore sequencing.

**EPV488:** EPV487 receiver strain; natural transformation with pEP102 for $P_{matP}$::*matP-3xFlag-aad1*, control PCR 3128+3162.

**EPV496:** JVV024 receiver strain; natural transformation with pJMD18 for $P_{matP}$::*matP-3xFlag-Sh ble*, control PCR 3128+3162.

**EPV498:** EPV487 receiver strain; natural transformation with pEP104 for *lacI-RFPT-YGFP-*$parB^{pMT1}$*-aad1* at *lacZ*, control PCR 600+2300; natural transformation with gDNA for *lacO* array-*aph* at DII, control PCR 1514+1764; natural transformation with pEG424 for $parS^{pMT1}$ at inverted DI, control PCR 503+2056.

**EPV499:** EPV50 receiver strain; natural transformation with pEG419 for $P_{matP}$::*matP*Δtetra-*3xFlag-aad1*; control PCR 3128+3162.

**EPV501:** EPV50 receiver strain; natural transformation with pEG420 for $P_{matP}$::*matP*Δdimer+tetra-*3xFlag-aad1*; control PCR 3128+3858.

**JMDV69:** EPV50 receiver strain; natural transformation with pJMD18 for $P_{matP}$::*matP-3xFlag-Sh ble*, control PCR 3128+3162.

**JMDV163:** MCH1 receiver strain; natural transformation with pJMD18 for $P_{matP}$::*matP-3xFlag-Sh ble*, control PCR 3128+3162.

**JMDV182:** EPV390 receiver strain; natural transformation with pJMD18 for $P_{matP}$::*matP-3xFlag-Sh ble*, control PCR 3128+3162.

**JVV013:** EPV487 receiver strain; natural transformation with pBJ3(3)1 for $P_{BAD}$::*xerC*; control PCR 595+1036.

**JVV018:** JVV013 receiver strain; natural transformation with pGD243 for *matP::bla*, control PCR 1857+1862.

**JVV024:** EPV50 receiver strain; natural transformation with pEG421 to integrate *vc1488* with its 2 *matS* sites between *vca0560* and *vca0561*, control PCR 3118+3555; natural transformation with pEG423 to delete *vc1488*, control PCR 3802+3791.

**STRAINS USED IN EACH FIGURE PANEL**

**Figure 1.** JMDV69.

**Figure 2.** B057 (WT) and C207 (Δ*matP*).

**Figure 3. (a, b, c)** L5I: WT (EGV339, EGV475), Δ*matP* (EGV331, EGV479); L6I: WT (EGV381, EGV477), Δ*matP* (EGV392, EGV481); DI: WT (EGV114, EGV324, EGV360, GDV548, EGV468, EGV475, EGV476, EGV477, EGV478), Δ*matP* (EGV326, EGV479, EGV480, EGV481, EGV482); R6I: WT (EGV382, EGV478), Δ*matP* (EGV393, EGV482); R5I: WT (EGV340, EGV476), Δ*matP* (EGV332, EGV480); R4I: WT (EGV341), Δ*matP* (EGV333); R2I: WT (EGV338), Δ*matP* (EGV330); OI: WT (EGV324, EGV325, EGV338, EGV339, EGV340, EGV341,EGV342, EGV343, EGV346, EGV381, EGV382), Δ*matP* (EGV326, EGV327, EGV330, EGV331, EGV332, EGV333, EGV334, EGV335, EGV347, EGV392, EGV393); OII: WT (EGV346), Δ*matP* (EGV347); L2II: WT (EGV342, EGV494), Δ*matP* (EGV334, EGV495); DII: WT (EGV325, EGV369, GDV548, EGV468, EGV494, EGV498), Δ*matP* (EGV327, EGV495, EGV499); R3II: WT (EGV343, EGV498), Δ*matP* (EGV335, EGV499). **(d)** WT (EEV29), Δ*matP* (EEV94). **(e)** WT (GDV28), Δ*matP* (EEV92).

**Figure 4. (b)** JMDV163. **(c ,d, e)** L5I: MCH1 (EGV225), MCH1_Δ*matP* (EGV241); L6I: MCH1 (EGV267), MCH1_Δ*matP* (EGV449); R1II: MCH1 (EGV313), MCH1_Δ*matP* (EGV450); DII: MCH1 (EGV224, EGV225, EGV226, EGV229, EGV232, EGV233, EGV234, EGV266, EGV267, EGV313), MCH1_Δ*matP* (EGV241, EGV242, EGV448, EGV449, EGV450, EGV451); L1II: MCH1 (EGV229); OII: MCH1 (EGV232), MCH1_Δ*matP* (EGV242); R5I: MCH1 (EGV226); R3I: MCH1 (EGV234); R1I: MCH1 (EGV233); OI: MCH1 (EGV224), MCH1_Δ*matP*

(EGV451). **(f)** MCH1 (EEV104), MCH1_Δ*matP* (EEV176). **(g)** MCH1 (EGV637), MCH1_Δ*matP* (EGV638).

**Figure 5. (b)** EPV496. **(c)** JVV018. **(d,e)** N16961 (EGV360, GDV548, EGV468), JJV024 (EGV503), EPV487 (EPV498). **(g)** EPV488**. (h)** JVV013.

**Figure 6. (a)** WT (EGV324, EGV325), Δ*matP* (EGV326, EGV327), *matP* Δ*dimer* (EGV452, EGV456), *matP* Δ*tetra* (EGV454, EGV455), Δ*zapA* (EGV336, EGV337), Δ*zapB* (EGV328, EGV329). **(b)** WT (EGV57), Δ*matP* (EGV353), *matP* Δ*dimer* (EGV632), *matP* Δ*tetra* (EGV633), Δ*zapA* (EGV426), Δ*zapB* (EGV425). **(c)** *ftsZ84* (EGV363), *ftsZ84* Δ*matP* (EGV365).

**Supplementary Figure 1. (b)** JMDV69 (MatP-Flag) and EPV501 (MatPΔdimer-Flag). **(c)** JMDV69.

**Supplementary Figure 2.** JMDV69.

**Supplementary Figure 3.** As in Figure 2.

**Supplementary Figure 4.** As indicated in the figure.

**Supplementary Figure 5. (a, b, c)** as in Figure 3. **(d)** GDV28.

**Supplementary Figure 6. (a)** as in Figure 4b. **(b, c)** as indicated in the figure. **(d)** as in Figure 4c. **(e)** as in Figure 4f. **(f)** as in Figure 4g.

**Supplementary Figure 7. (a)** as in Figure 5c. **(b)** as indicated in the figure. **(c)** as in Figure 5d. **(d)** as in Figure 5h. **(e)** EPV487 (JJV013), EPV487 Δ*matP* (JVV018).

**Supplementary Figure 8. (a)** EPV499. **(b)** as indicated in the figure.

**Supplementary Figure 9. (a, b)** as in figure 6b. **(c)** JMDV182.
